# Stable colony stimulating factor 1 fusion protein treatment increases HSC pool and enhances their mobilisation in mice

**DOI:** 10.1101/2020.09.13.292227

**Authors:** Simranpreet Kaur, Anuj Sehgal, Andy C. Wu, Susan M Millard, Lena Batoon, Michelle Ferrari-Cestari, Jean-Pierre Levesque, David A. Hume, Liza J. Raggatt, Allison R. Pettit

**Affiliations:** Mater Research Institute-The University of Queensland, Faculty of Medicine, Translational Research Institute, Woolloongabba, 4102, Australia

## Abstract

Prior chemotherapy and/or underlying morbidity commonly leads to poor mobilisation of hematopoietic stem cells (HSC) for transplantation in cancer patients. Increasing the number of available HSC prior to mobilisation is a potential strategy to overcome this deficiency. Resident bone marrow (BM) macrophages are essential for maintenance of niches that support HSC and enable engraftment in transplant recipients. Here we examined potential of donor treatment with colony stimulating factor-1 (CSF1) to modify the BM niche and expand the potential HSC pool for autologous transplantation. We administrated CSF1 Fc fusion protein (CSF1-Fc) to naive C57Bl/6 mice and assessed the impacts on HSC number and function and overall haematopoiesis. Outcomes were assessed by *in situ* imaging and *ex vivo* flow cytometry with functional validation by colony formation and competitive transplantation assay. CSF1-Fc treatment caused a transient expansion of monocyte-macrophage cells within BM and spleen at the expense of BM B lymphopoiesis and hematopoietic stem and progenitor cell (HSPC) homeostasis. During the recovery phase after cessation of CSF1-Fc treatment, normalisation of haematopoiesis was accompanied by an increase in the total available HSPC pool. In the spleen, increased HSC was associated with expression of the BM niche marker CD169 in red pulp macrophages. Pre-treatment with CSF1-Fc increased the number and reconstitution potential of HSPC in blood following a HSC mobilising regimen of granulocyte colony stimulating factor (G-CSF) treatment. These results indicate that CSF1-Fc conditioning could represent a therapeutic strategy to overcome poor HSC mobilisation and subsequently improve autologous or heterologous HSC transplantation outcomes.

**Key points:** 1) Recovery from Fc-modified colony stimulating factor-1 (CSF1-Fc) treatment was accompanied by an increase in total haematopoietic stem cells. 2) Pre-conditioning with CSF1-Fc increased the reconstitution potential of blood after haematopoietic stem cell mobilisation.

## Introduction

Mobilisation of hematopoietic stem and progenitor cells (HSPC) into peripheral blood is used to enable collection of enriched hematopoietic stem cells (HSC) for transplantation. This procedure is a successful approach used to treat a broad range of immune and haematological malignancies and deficiencies. In up to 40% of patients referred for autologous transplant, insufficient numbers of HSPC are mobilised due to underlying morbidity or prior treatment impacts on the HSPC pool^1^. This deficiency can preclude HSC transplant in these poor mobilisers leaving no other effective treatment options^2^. Development of approaches to achieve HSC expansion prior to mobilization would address this treatment gap^3^.

HSC reside in specific locations called niches in the bone marrow (BM) and BM resident macrophages are an integral component of HSC niches. Macrophage depletion *in vivo* is sufficient to drive mobilisation of HSC to blood^4–6^ and prevent successful re-establishment of the HSC niche after total body irradiation^7^. Granulocyte colony stimulating factor (G-CSF) was one of the first growth factors used to mobilise HSC^8^ and remains the main compound to elicit HSPC mobilisation in donors and patients^9^. G-CSF triggers a complex array of mechanisms affecting HSC niche cellular and structural components^10^, including direct effects on BM macrophages to elicit stem cell mobilisation^4,11^.

Colony stimulating factor-1 (CSF1) is required for the differentiation, survival and proliferation of tissue resident macrophages^12,13^. Although clinical trial of CSF1 as an adjunct therapy for patients receiving HSC transplantation showed some benefit^14^, it has not progressed into mainstream clinical care. Recombinant CSF1 is rapidly cleared by the kidneys and the clinical use of CSF1 requires repeat high dose/continuous infusion (reviewed in^12^), making clinical use and pre-clinical studies impractical and/or cost-prohibitive. To address CSF1 therapeutic limitations, Gow *et. al*. engineered a pig CSF1 molecule conjugated to the Fc (CH-3) region of pig immunoglobulin IgG1a that greatly increased its circulating half-life^15^. Pig CSF1 is equally active on mouse and human macrophages^16^ and short term daily CSF1-Fc injection was well tolerated and effectively increased murine blood monocyte and tissue resident macrophage populations, including those in BM, in mice^15^, rats^17^ and pigs^18^. However, the improved drug qualities of CSF1-Fc are associated with supraphysiologic pharmacokinetic properties^12,15,19^. This, as well as the resultant expansion of resident macrophages induced by treatment, could have complex consequences on haematopoiesis in BM and spleen that have not been investigated.

We hypothesised that CSF1-Fc treatment has direct and indirect impacts on the HSPC compartment that might be harnessed to improve HSC transplantation outcomes. Herein, we studied the temporal profile of the effects of an acute CSF1-Fc treatment regimen on resident macrophages, the HSPC pool and developing as well as mature leucocytes and erythroid cells. While the treatment initially depleted HSPC within the BM and triggered extramedullary haematopoiesis, the recovery phase following treatment was associated with an increase in total HSC that could be mobilised using subsequent G-CSF administration. We suggest that CSF1-Fc has potential utility in conditioning for more effective stem cell mobilisation.

## Methods

### Animals

All procedures complied with the Australian Code of Practice for the Care and Use of Animals for Scientific Purposes and were approved by The University of Queensland (UQ) Health Sciences Ethics Committee. C57BL/6 (CD45.2^+^) mice were sourced from Australian Resources Centre. Congenic B6.SJL-*Ptprc*^*a*^ *Pep*^*c*^/BoyJ mice (B6.SJL CD45.1^+^), competitor transgenic red fluorescent protein (RFP) mice were generated by Professor Patrick Tam (Children’s Medical Research Institute, Sydney, Australia) and derived from TgN (ACTB-DSRed.T3) Nagy ES cells^20^ and C57BL/6-Tg(UBC-GFP)30^Scha/J^ green fluorescent protein (GFP) reporter mice were supplied from in-house breeding colonies. All mice used were 10-week old females housed under specific pathogen-free conditions.

### *In vivo* studies

C57BL/6 mice were treated daily with either CSF1-Fc (1 mg/kg) or saline subcutaneously for 4 consecutive days as previously described^15^. Mice were sacrificed at 7 days and 14 days post-first saline or CSF1-Fc injection and peripheral blood^21^, spleen^22^, liver^23^ and femoral bone^24^ were harvested for flow cytometry assessment, immunohistochemistry and colony forming unit (CFU) assays^25^. HSC mobilization was performed 14 days post-first CSF1-Fc treatment by 3 consecutive bi-daily intraperitoneal injections of saline or G-CSF (125 µg/kg; Filgrastim, Amgen, Thousand Oaks, CA) with the same tissues collected the day following the last G-CSF injection.

### Competitive transplant assays

Competitive grafts were generated using 2 × 10^5^ BM from C57BL/6 donor mice treated with saline or CSF1-Fc (as per above) and equal numbers of BM cells from naïve UBC-GFP ‘competitor’ mice as described^26^. The BM cells were injected intravenously into lethally irradiated 10-week old C57BL/6 recipient female mice. In another competitive transplant assay, C57BL/6 donor mice were treated with combination of CSF1-Fc or saline (as per above) plus G-CSF and whole heparinized blood was collected via cardiac puncture. 20 µl of whole blood was then mixed with 2 × 10^5^ BM competitor cells from naïve RFP mice to generate the competitive graft^26^. The competitive grafts were intravenously injected into lethally irradiated 10-week-old B6.SJL CD45.1^+^ recipient mice. Blood was collected from secondary recipients at 8-, 12- and 16-week post-transplant to quantify chimerism. Content in repopulation units (RU) was calculated for each individual recipient mouse according to CD45.2^+^ donor blood chimerism at 16 weeks post-transplantation using the following formula: 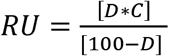 where D is the percentage of donor CD45.2^+^ B and myeloid cells and C is the number of competing CD45.1^+^ BM RUs co-transplanted with the donor cells (C=2 for 200 000 competing BM cells). RUs were then converted to per ml of blood^27^.

### Flow cytometry

Myeloid lineage^7,28^, HSPC ^7,29^, B cell subsets^30,31^ and red blood cell maturation^29,32,33^ phenotyping were performed as detailed in Supplementary Table 1. Cell acquisition was performed on Beckman Coulter’s CyAn™ ADP Analyser (Beckman Coulter, USA), Cytoflex Analyser (Beckman Coulter, USA) or BD LSRFortessa™ X-20 (BD Biosciences, USA) for panels specified in Supplementary Table 1. Data analysis was performed using the FlowJo software (Tree Star Data Analysis Software, Ashland, OR).

### Immunohistochemistry and immunofluorescence

Five µm sections of left hind limb and spleen, sampled from each block at 3 sectional depths 100 µm apart, were deparaffinized and rehydrated prior to staining with rat anti-F4/80 monoclonal antibody (Abcam, UK) or isotype control as previously described^34^. Sections were scanned at 40X magnification using Olympus VS120 slide scanner (Olympus, Japan) and analyzed for percent area of chromogen staining in the entire section using Visiopharm VIS 2017.2 Image analysis software (Hørsholm, Denmark). Representative images were collected on an Olympus Bx50 microscope and Cell Sens standard software 7.1 (Olympus, Tokyo, Japan). For immunofluorescence, spleens were snap frozen in liquid nitrogen and embedded in Optimal Cutting Temperature (OCT; VWR, UK). Frozen sections (5 μm) were stained with rat anti-F4/80 (Abcam, UK) followed by species-specific secondary antibody coupled to Alexa Fluor (AF) 647 (Invitrogen, USA). Additional staining with anti-mouse CD45R (B220)-AF488 (Biolegend, USA), anti-mouse CD169-AF594 (Biolegend, USA) or anti-mouse CD3ε-AF647 (Biolegend, USA) was performed. Immunofluorescence images were acquired on the Upright Motorized Olympus BX63 Upright Epifluorescence Microscope (Olympus Life Science, Australia). For morphometric analysis, digital microscopy images were analyzed using ImageJ software (http://rsb.info.nih.gov/ij/) as previously described^35^. Briefly, all images were coded and assessed blindly. Background intensity thresholds were applied using an ImageJ macro which measures pixel intensity across all immunostained and non-stained areas of the images.

### Statistical Analysis

Details of all group/sample sizes and experimental repeats are provided in figure legends. Unless indicated otherwise, data are mean ± standard deviation (SD). Statistical analysis was performed using one-way ANOVA with Tukey’s multiple comparison test. In instances where there was evidence of non-normality identified by the Kolmogorov–Smirnov test, data were analysed by a Mann–Whitney U-test. Values of P < 0.05 were accepted as significant.

## Results

### Acute CSF1-Fc treatment expands BM monocytes and resident macrophages

CSF1-Fc treatment was previously demonstrated to increase F4/80^+^ and Gr-1^+^ BM cells at day 5 after 4 consecutive daily treatments^15^. However, sustained impacts and resolution kinetics of CSF1-Fc treatment on BM myeloid and broader impacts on other hematopoietic lineages were not mapped. Given CSF1-Fc remains substantially elevated in circulation for 72 hr post-delivery, peak treatment effects could be delayed by at least 3 days post-treatment cessation. Consequently, CSF1-Fc was administered to female C57BL/6 mice daily for 4 days as described^15^ and BM, spleen and blood were examined at 7 and 14 days post-initiation of treatment (Fig. 1a). F4/80^+^ resident BM macrophages and monocytes were present throughout the BM in saline treated mice (Fig. 1b) including perivascular (closed arrows) and endosteal (open arrows) regions known to be enriched for HSC niches^36^. 7 days after initial CSF1-Fc treatment the frequency of F4/80^+^ cells and their apparent ramification in BM was appreciably increased (Fig. 1b) and could be quantified as a 1.8-fold expansion in F4/80 staining area (Fig. 1c). This F4/80^+^ macrophage/monocyte expansion was transient and returned to baseline levels by day 14 (Fig. 1b and c). Flow cytometry confirmed a relative increase in BM monocytes at day 7 which returned to baseline by day 14 (Fig. 1d; Supp. Fig. 1a). There was no change in BM granulocyte number compared with saline at either time point (Fig. 1e; Supp. Fig. 1b). BM B cell frequency was decreased at day 7 and rebounded to supraphysiologic levels at day 14 (Fig. 1f; Supp Fig. 2a). BM T cells were not immediately affected by treatment but were modestly increased at day 14 (Fig. 1g; Supp Fig. 2a). Overall, CSF1-Fc treatment stimulated a robust but transient expansion of BM monocytes and macrophages and while it disrupted B lymphopoiesis, this was also transient and corrected by a rebound expansion of BM lymphocytes.

**Fig. 1.**
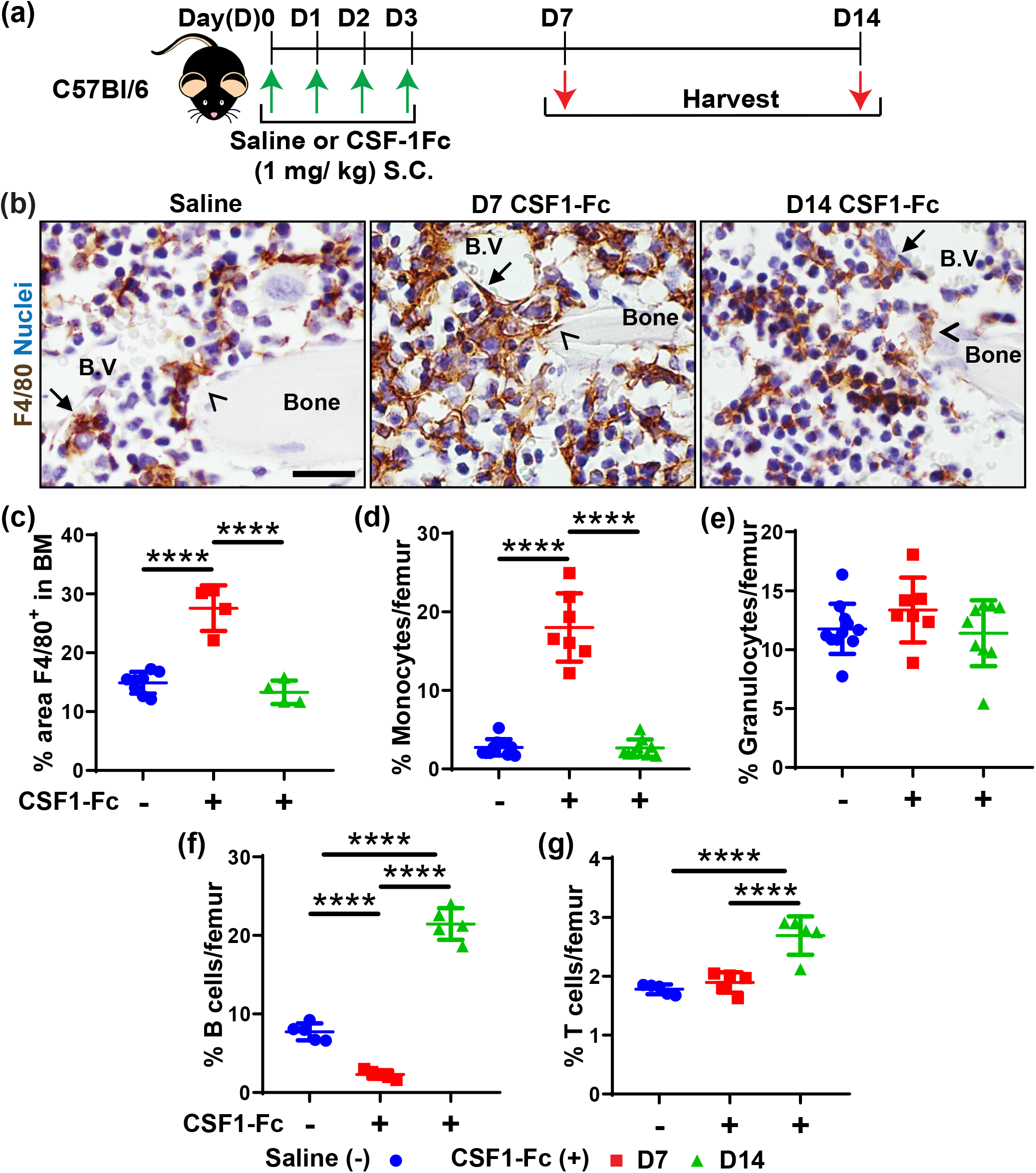
CSF1-Fc treatment induced significant expansion of BM resident macrophages. (**a)** Schematic of CSF1-Fc treatment regimen in C57BL/6 mice (D, day; S.C., subcutaneous). Tissues were harvested at 7 (D7) and 14 days (D14) post-first CSF1-Fc injection. **(b)** F4/80 immunohistochemistry (brown) in femoral BM sections of mice treated with saline (left panel) or CSF1-Fc at D7 (middle panel) and D14 (right panel). Closed arrows indicate perivascular macrophages and arrowheads highlight endosteal macrophages. Sections were counterstained with haematoxylin (blue) and taken at 600X magnification. Scale bar = 20 µm. (**c)** Quantification of percent area of F4/80 staining in the femur of saline (blue circles, pooled D7 and D14 samples) and CSF1-Fc treated mice at D7 (red squares) and D14 (green triangles) post-first injection. **(d-g)** Flow cytometry analysis to determine percentage frequency of **(d)** F4/80^+^Ly6G^neg^VCAM^neg^CD115^+^CD11b^+^ monocytes, **(e)** CD11b^+^Ly6G^+^ granulocytes, **(f)** CD11b^neg^CD3^neg^B220^+^ B cells and **(g)** CD11b^neg^B220^neg^CD3^+^ T cells in BM of saline controls or CSF1-Fc treated mice. Flow cytometry representative raw data and gating provided in Supp. Fig. 1 and 2. Each data point represents a separate mouse and bars are mean ± SD. Statistical analysis was performed using one-way ANOVA Tukey’s multiple comparison test where ****p<0.0001.

**Fig. 2.**
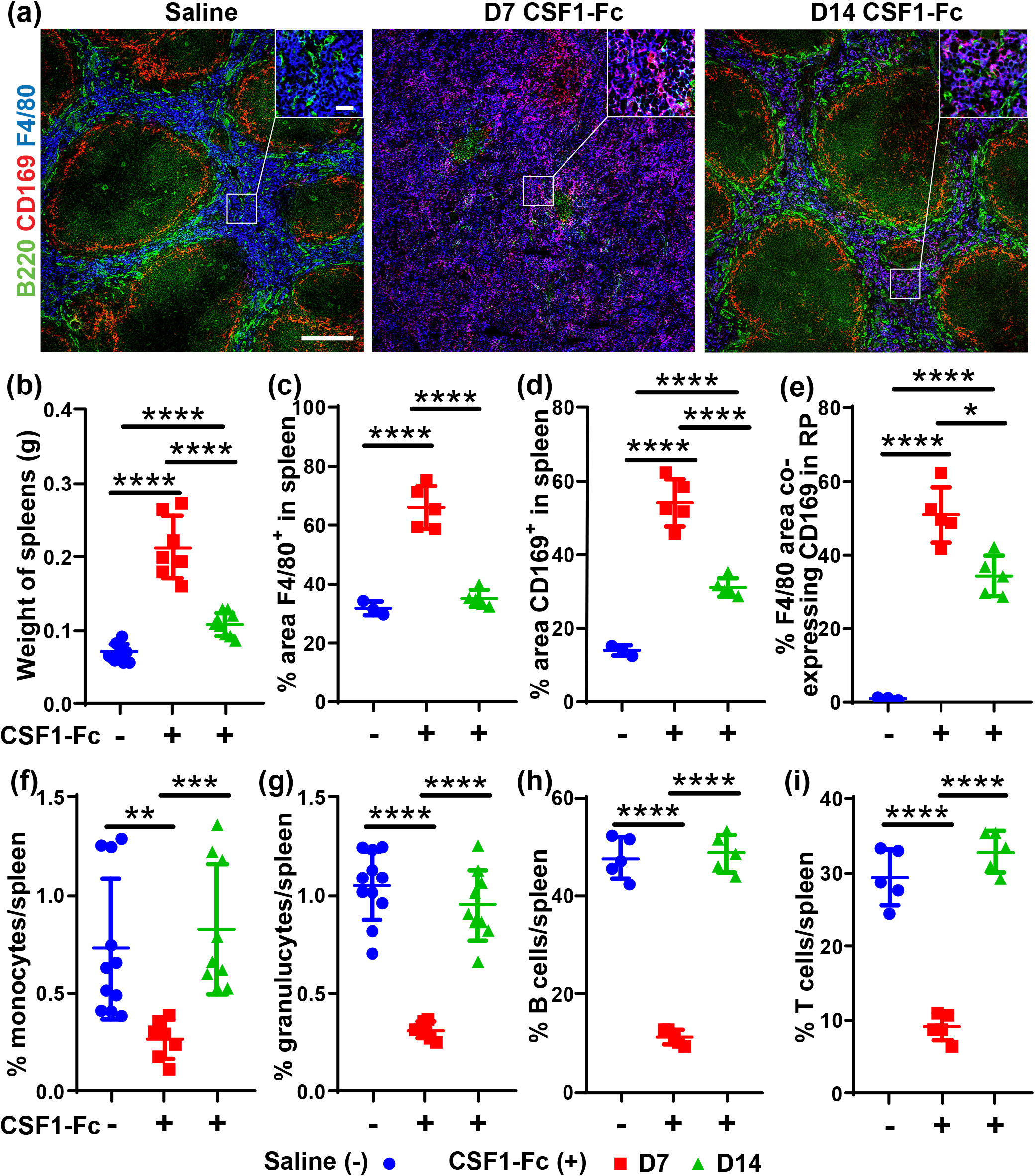
CSF1-Fc treatment transiently increased splenic macrophages, disrupted splenic architecture, and altered red pulp macrophage phenotype. **(a)** Immunofluorescence labelling of F4/80 (blue), CD169 (red) and B220 (green) expression in spleen sections of mice treated with saline (left panel) or CSF1-Fc and assessed at D7 (middle panel) and D14 (right panel). Magnification = 100X; scale bar = 100 µm. Inset magnification = 600X; scale bar = 20 µm. **(b)** Spleen weights in saline controls or CSF1-Fc treated mice at the D7 and D14 time points. **(c-e)** Morphometric analysis of percent areas of **(c)** F4/80 immunolabelling, **(d)** CD169 immunolabelling and **(e)** F4/80 area co-labelled with CD169. **(f-i)** Flow cytometry analysis of percentage frequency of **(f)** F4/80^+^Ly6G^neg^VCAM^neg^CD115^+^CD11b^+^ monocytes, **(g)** CD11b^+^Ly6G^+^ granulocytes, **(h)** CD11b^neg^CD3^neg^B220^+^ B cells and **(i)** CD11b^neg^B220^neg^CD3^+^ T cells in spleens of saline controls or CSF1-Fc treated mice at both time points. Each data point represents a separate mouse and bars are mean ± SD. Statistical analysis was performed using one-way ANOVA Tukey’s multiple comparison test where ****p<0.0001, **p<0.01 and *p<0.05, n = 3 to 11 mice/group. Kolmogorov–Smirnov test revealed non-normality for data in graph **(f)** therefore dictating use of a Mann–Whitney U-test. Data for the saline control samples from D7 and D14 were pooled together in the graphical representations.

### Prolonged elevation of splenic resident macrophages post-CSF1-Fc treatment

Gow *et. al*. previously reported an increase in spleen weight at day 5 after the same CSF1-Fc treatment regimen in transgenic MacGreen mice. This was associated with increased number of splenic GFP^+^ myeloid cells as well as increased area of the red pulp and marginal zone^15^. *In situ* assessment of the prolonged effects of CSF1-Fc treatment on splenic myeloid populations revealed overt disruption of gross splenic morphology at day 7 after CSF1-Fc treatment in non-transgenic mice (Fig. 2). There was dramatic decrease in both B220^+^ B cell (Fig. 2a) and CD3^+^ T cell (Supp. Fig. 3a and b) staining with minimal identifiable white pulp on standard histology (not shown). There was also a complete loss of CD169^+^F4/80^low/-37^ marginal zone metallophilic macrophages (Fig. 2a) and a distinguishable marginal zone. Disrupted splenic architecture was accompanied by sustained splenomegaly with spleen weight increased 3-fold 7 days after CSF1-Fc injection and remained 1.4-fold larger at 14 days post initial treatment (Fig. 2b). There was a substantial increase in F4/80^+^ macrophage stained area within red pulp at day 7 post-treatment (Fig. 2a and c). In normal spleen, red pulp macrophages do not express CD169^37,38^ and this was confirmed in saline-treated mice (Fig. 2a and e). Notably, at day 7 in CSF1-Fc treated mice, the majority of red pulp macrophages in the expanded spleens of CSF1-Fc treated mice acquired detectable CD169 expression (Fig. 2a, d and e).

**Fig. 3.**
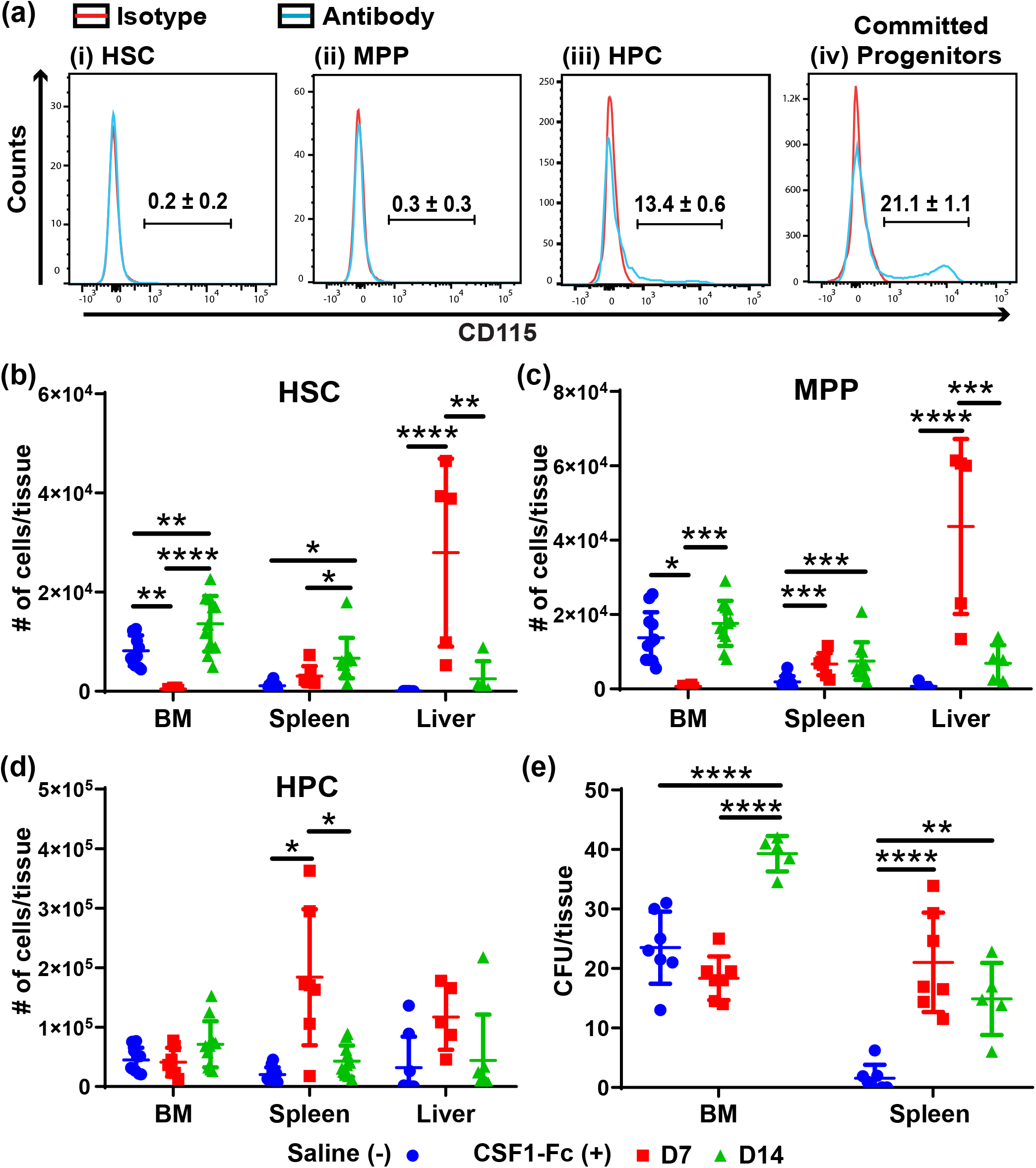
CSF1-Fc treatment is associated with a delayed increased HSC and MPP in BM and spleen. (**a)** Representative flow cytometry histograms of C57BL/6 mouse BM expression of CD115 in (i) HSC, (ii) MPP, (iii) HPC and (iv) committed progenitors. Population gating strategies are exemplified in Supp Figs. 4 and 6. The histograms show antibody staining (blue lines) compared to appropriate isotype staining (red lines). **(b-d)** Flow cytometry analysis to determine the number of **(b)** HSC, **(c)** MPP and **(d)** HPC in BM, spleen or liver of C57BL/6 mice treated with saline (blue circles) or CSF1-Fc at D7 (red squares) or D14 (green triangles) post-first CSF1-Fc injection. (**e)** Quantification of CFU in the BM (assay using single femur only) and spleen of saline or CSF1-Fc treated mice at both time points. Each data point represents a separate mouse and bars are mean ± SD. Statistical analysis was performed using one-way ANOVA Tukey’s multiple comparison test where ****p<0.0001, ***p<0.0005, **p<0.01 and *p<0.05, n = 4 to 11 mice/group. Kolmogorov–Smirnov test revealed non-normality for liver data in graphs **(b)** and **(c)** therefore dictating use of a Mann–Whitney U-test. Data for the saline control samples from D7 and D14 time points were pooled together in the graphical representations.

While the gross disorganisation of the spleen resolved by day 14, the altered F4/80^+^CD169^+^ red pulp macrophage phenotype persisted (Fig. 2a and 2e). Additionally, CD169^+^ macrophages were also detected within the CD3^+^ T cell zones of the white pulp in CSF1-Fc treated mice in contrast to saline treated controls (Supp. Fig. 3a and c). Flow cytometry showed a significant decrease in relative abundance of splenic monocytes (2.8-fold), granulocyte (3.3-fold), B cells (4.2-fold) and T cells (3.2-fold) at day 7 in CSF1-Fc treated mice which resolved by day 14 (Fig. 2f – 2i). Overall, CSF1-Fc treatment temporarily disrupted splenic architecture, characterised by expansion of red pulp macrophages at the expense of other mature leukocytes, but induced a prolonged increase in spleen size accompanied by a persistent alteration in red pulp macrophage phenotype.

### CSF1-Fc treatment transiently increased HSPC populations in BM, spleen and liver

CD169 is expressed by HSC^5,7^ and erythroid^39,40^ niche macrophages in the BM. Given both the sustained increase in spleen size and shift in splenic red pulp macrophage phenotype to express CD169^5,7^, we assessed whether CSF1-Fc treatment induced extramedullary haematopoiesis and/or impacted HSPC populations in BM, spleen and liver at 7 and 14 days post treatment. HSPC functional subsets were subdivided using signalling lymphocyte activation molecule (SLAM) marker^41^ expression in lineage negative, c-Kit^+^ and Sca1^+^ cells (LSK): CD48^neg^CD150^+^ self-renewing HSC, CD48^neg^CD150^neg^ non-self-renewing multipotent progenitors (MPP) and CD48^+^ hematopoietic progenitor cells (HPC). Committed progenitor cells were gated as lineage negative, c-Kit^+^ and Sca1^neg^ cells (Supp. Fig 4). Single cell transcriptional profiling indicates *Csf1r* mRNA is expressed in predominantly committed progenitors^42^ but mRNA expression can occur prior to detectable cell surface CSF1R protein^43,44^ and in granulocytes *Csfr1* mRNA does not get translated *in vivo*^45^. Consistent with these earlier findings, HSC and MPP did not express detectable CSF1R, whereas it was detected on a small subpopulation of HPC and committed progenitors (Fig. 3a). Nevertheless, CSF1-Fc treatment induced dynamic changes in phenotypic BM HSC and MPP. Both populations were significantly decreased at 7 days post treatment (Fig. 3b and 3c). The MPP returned to normal (Fig. 3c) but the HSC numbers rebounded to 1.7-fold above baseline at day 14 (Fig. 3b) w. By contrast, HPC were unchanged by CSF1-Fc treatment (Fig. 3d).

**Fig. 4.**
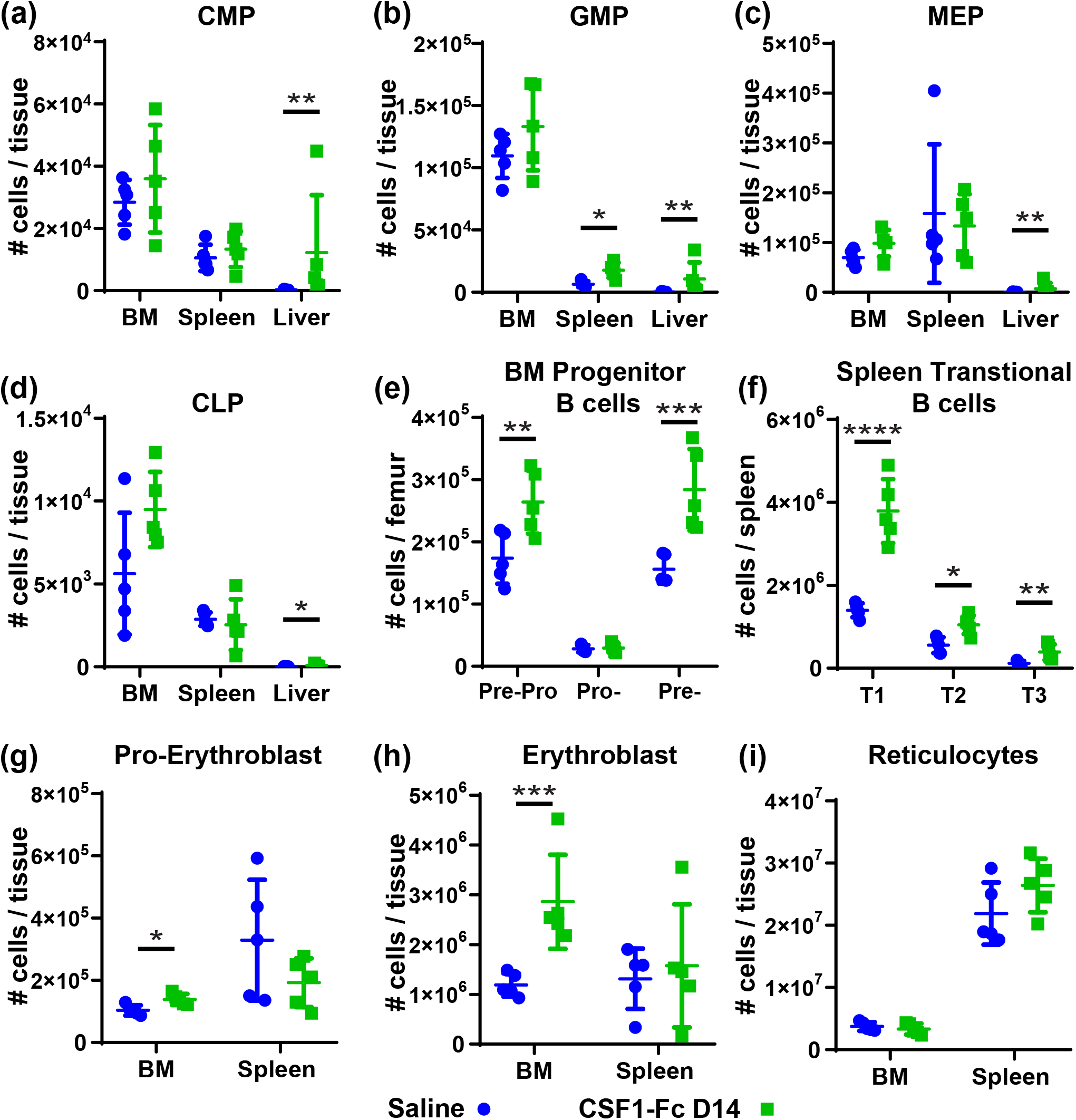
CSF1-Fc treatment recovery was not associated with splenic extramedullary haematopoiesis. **(a-d)** Flow cytometry analysis to determine cell number of **(a)** CMP, **(b)** GMP, **(c)** MEP cells and **(d)** CLP cells in BM, spleen and liver of C57BL/6 mice treated with saline (blue dots) or CSF1-Fc at D14 (green triangles) post-first CSF1-Fc injection. Population gating strategies are exemplified in Supp Fig. 6. **(e)** Flow cytometry analysis of BM B cell progenitor subsets at 3 developmental stages: Pre-Pro B cells, Pro-B cells and Pre-B cells in BM of C57BL/6 mice treated as above. Population gating strategies are exemplified in Supp Fig. 2. **(f)** Flow cytometry analysis of splenic transitional (T1-3) B cell maturation: CD93^+^CD23^neg^IgM^+^ T1 B cells, CD93^+^CD23^+^IgM^+^ T2 B cells and CD93^+^CD23^+^IgM^neg^ T3 B cells. Population gating strategies are exemplified in Supp. Fig 2. **(g-i)** Flow cytometry analysis of **(g)** Ho^+^Ter119^low^CD71^+^ pro-erythroblasts, **(h)** Ho^+^Ter119^+^CD71^+^ erythroblasts and **(i)** Ho^neg^Ter119^+^ reticulocytes BM (quantification in single femur only) and spleen of C57BL/6 mice treated as above. Each data point represents a separate mouse and bars are mean ± SD. Statistical analysis was performed using one-way ANOVA Tukey’s multiple comparison test where ****p<0.0001, ***p<0.0005, **p<0.01 and *p<0.05.

The CSF1-Fc-induced reduction of detectable BM HSC and MPP at day 7 could be a consequence of direct or indirect regulation of the key marker c-Kit^21,46^. To confirm that these cell populations were genuinely altered by CSF1-Fc treatment, colony-forming unit (CFU; Fig. 3e) and competitive transplantation assays were performed (Supp. Fig. 5a). There was no significant difference in CFU potential in BM from saline or day 7 post CSF1-Fc treated animals (Fig. 3e). However, competitive transplant confirmed loss of repopulation potential compared to saline-treated donor BM (Supp. Fig. 5b).

**Fig. 5.**
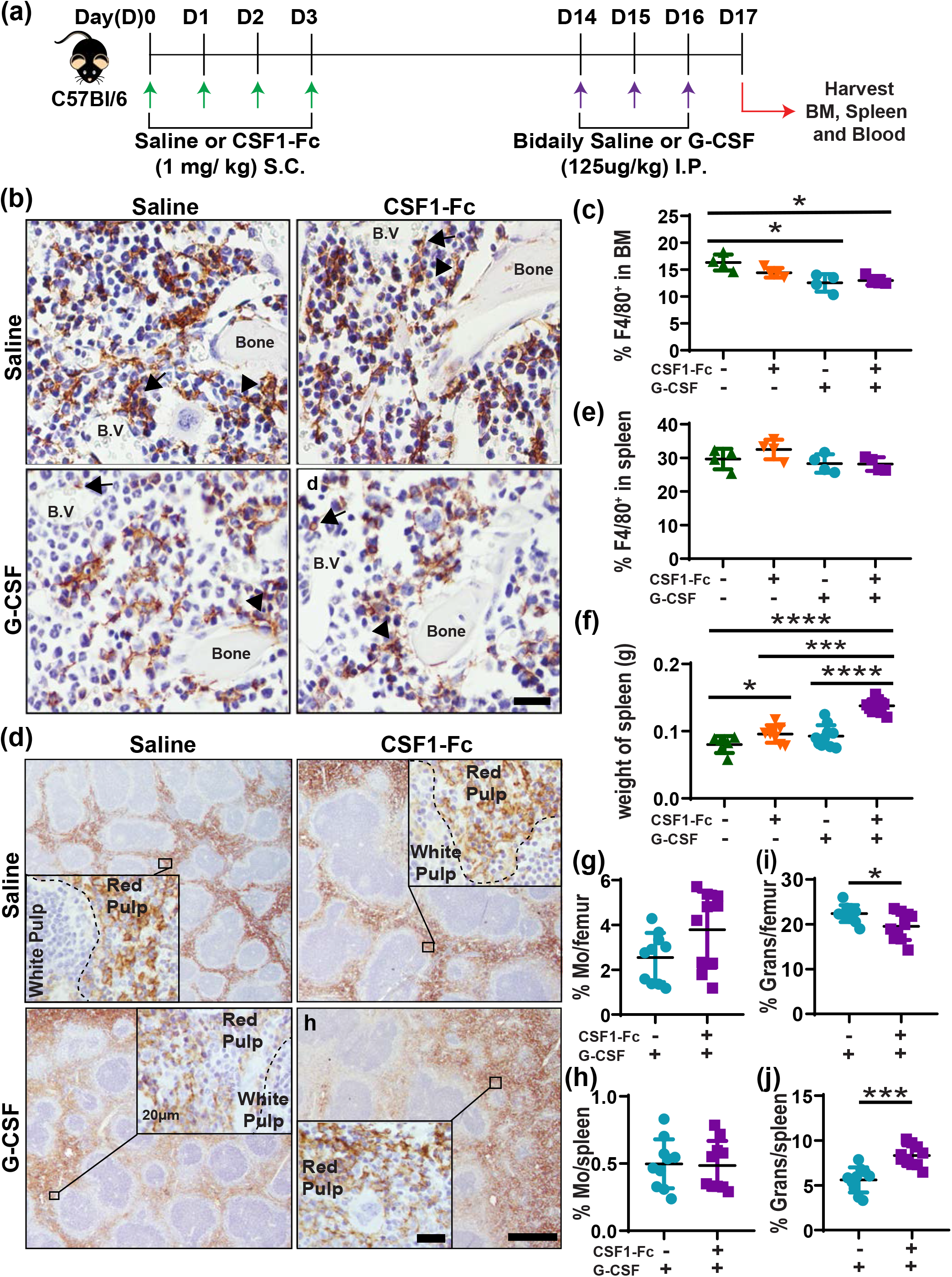
Combination treatment with CSF1-Fc + G-CSF had only modest cumulative impact on BM and spleen myeloid cells. (**a)** Schematic of tandem CSF1-Fc plus G-CSF treatment regimen to C57BL/6 non-transgenic mice. Briefly, mice were divided into 4 treatments groups: (1) once daily saline for 4 days followed by bi-daily saline treatment initiated 14 days later (saline + saline); (2) once daily CSF1-Fc for 4 days followed by bi-daily saline treatment initiated 14 days later (CSF1-Fc + saline); (3) once daily saline for 4 days followed by bi-daily G-CSF treatment initiated 14 days later (saline + G-CSF); and (4) once daily CSF1-Fc for 4 days followed by bi-daily G-CSF treatment initiated 14 days later (CSF1-Fc + G-CSF). Tissues were harvested 17 days post-first CSF1-Fc injection. **(b)** Representative immunohistochemistry anti-F4/80 staining (brown) in femoral sections of mice treated with saline + saline (top left panel), CSF1-Fc + saline (top right panel), saline + G-CSF (bottom left panel) and CSF1-Fc + G-CSF (bottom right panel). F4/80^+^ BM resident macrophages (brown) are lining the endosteal (arrowheads) and perivascular (arrow) regions of the BM. Sections were counterstained with haematoxylin (blue) and taken at 600X magnification; scale bar = 20 µm. **(c)** Quantification of percent area of F4/80 staining in the femur of mice treated as above. **(d)** Representative immunohistochemistry anti-F4/80 staining (brown) in splenic sections of mice treated as above. Section were counterstained with haematoxylin (blue) and taken at 100X magnification; scale bar = 500 µm. Inset at 600X magnification; scale bar = 20 µm. **(e)** Quantification of percent area of F4/80 staining in the spleen of mice treated as above. **(f)** Weights of spleen of mice treated as above. **(g-j)** Flow cytometry analysis of the percentage of F4/80^+^Ly6G^neg^VCAM^+^CD115^+^CD11b^+^ monocytes (Mo) in BM **(g)** and spleen **(h)** or CD11b^+^Ly6G^+^ granulocytes (Grans) in BM **(i)** and spleen **(j)** of mice treated with either saline + G-CSF and CSF1-Fc + G-CSF. Each data point represents a separate mouse and bars are mean ± SD. Statistical analysis was performed using one-way ANOVA Tukey’s multiple comparison test and unpaired student t-test where ****p < 0.0001, ***p<0.001, *P<0.05.

Unexpectedly, CSF1-Fc treatment induced a sustained increase of both phenotypic HSC and MPP in the spleen with these populations remaining elevated at day 14 (Fig. 3b and 3c). Splenic HPC were substantially increased at day 7 but returned to normal by day 14 (Fig. 3d). The treatment also led to a transient increase in phenotypic HSC and MPP in the liver at 7 days post treatment which resolve by day 14 (Fig. 3b and c). No change was noted in liver HPC number (Fig. 3d). CSF1-Fc treatment did not induce HSPC mobilisation into blood at either time point (not shown). In the spleen, CFU activity aligned with the large and sustained increase in phenotypic HSPC driven by CSF1-Fc treatment (Fig. 3e). Overall, these data indicate that CSF1-Fc treatment produced a transient decrease in HSPC in BM at day 7 followed by a compensatory overshoot in the HSPC pool in BM and spleen at 14 days after initial treatment. To determine whether the increased splenic HSC pool observed at 14 days post-CSF1-Fc treatment was an indirect consequence of compensatory extramedullary haematopoiesis, myeloid progenitors^7,28^, lymphoid progenitors^29,30,47^, erythroblasts and reticulocytes^32,33^ were assessed in BM, spleen and liver at day 14 post initial CSF1-Fc or saline treatment. No effect on common myeloid progenitors (CMP) number was observed at this time point in either BM or spleen, whereas they were increased in the liver (Fig. 4a). Granulocyte macrophage progenitor (GMP) number was not impacted in BM but was elevated in spleen and liver (Fig. 4b) and this is likely attributable to sustained CSF1-Fc treatment impacts on myeloid lineage dynamics. Megakaryocyte erythroblast progenitors (MEP) and common lymphoid progenitors (CLP) were unchanged at day 14 in the BM and spleen but increased in the liver (Fig. 4c and d).

The rebound in B cell populations noted in Figure 1 and 2 in BM and spleen is likely attributable to significantly increased Pre-Pro B cells and Pre-B cell progenitor subsets in the BM (Fig. 4e) at day 14 with no change in the spleen (not shown), where progenitor B cells do not reside under homeostatic conditions^31,48^. All 3 transitional B cell populations that are normally located within spleen^30,31^ were increased at day 14 after CSF1-Fc treatment (Fig. 4f) indicative of systemic elevation of B lymphopoiesis. Erythropoiesis, as indicated by number of proerythroblasts (Fig. 4g) and erythroblasts (Fig. 4h), was elevated in BM but not in the spleen when compared to saline control at day 14 post-CSF1-Fc. Reticulocytes number was equivalent to baseline in both BM and spleen (Fig. 4i).

It should be noted that evidence of splenic extramedullary haematopoiesis, predominantly erythropoiesis, was evident at day 7 (not shown). However, the data presented in Fig. 4, combined with reestablishment of normal splenic morphology by day 14 (Fig. 2a; Supp. Fig. 3a), suggest that the increased splenic HSPC pool was not a consequence of sustained extramedullary haematopoiesis within this spleen during recovery from CSF1-Fc treatment. A more likely explanation consistent with induced expression of CD169 (Fig. 2a), is that the treatment has changed splenic capacity to support HSPC homing and/or temporary lodgement. On this basis, with appropriate timing, CSF1-Fc treatment has the potential to expand available quiescent HSPC for mobilisation.

### G-CSF-induced HSPC mobilisation efficiency is enhanced by pre-treatment with CSF1-Fc

To test CSF1-Fc as a potential donor conditioning regimen, mice were treated with CSF1-Fc for 4 days followed by a HSPC mobilising regimen of G-CSF initiated at day 14 (CSF1-Fc + G-CSF, Fig 5a). Consistent with a previous study^4^, G-CSF treatment reduced F4/80^+^ macrophage frequency in BM (Fig. 5b and c) but had no significant impact on F4/80^+^ macrophage frequency in spleen (Fig. 5d and e). These responses were not altered by CSF1-Fc conditioning. Spleen weights of CSF1-Fc + G-CSF treated mice were significantly higher when compared to the other 3 treatment groups (Fig. 5f). For simplicity, only G-CSF treated groups are shown in subsequent figures. Pre-CSF1-Fc treatment did not change monocyte frequency in BM (Fig. 5g) or spleen (Fig. 5h) of G-CSF treated mice. BM granulocyte frequency was significantly reduced (Fig. 5i) while splenic granulocyte frequency was significantly increased (Fig. 5j) by the combination therapy compare to G-CSF treatment alone. No significant change in HSC, MPP or HPC number in BM of G-CSF mobilised mice was induced by pre-treatment with CSF1-Fc (Fig. 6a-c). However, an increase in circulating phenotypic HSC (2.5-fold), MPP (2.5-fold) and HPC (3.8-fold) mobilised by G-CSF were each increased in the blood of CSF1-Fc pre-treated mice compared to controls (Fig. 6d-f). Similar relative increases in HSC, MPP and HPC were detected in spleen in mice pre-treated with CSF1-Fc (Fig. 6g-i). The effect of the pre-treatment on progenitor cell mobilisation was confirmed by CFU assays using BM, blood or spleen (Fig. 6j-l).

**Fig. 6.**
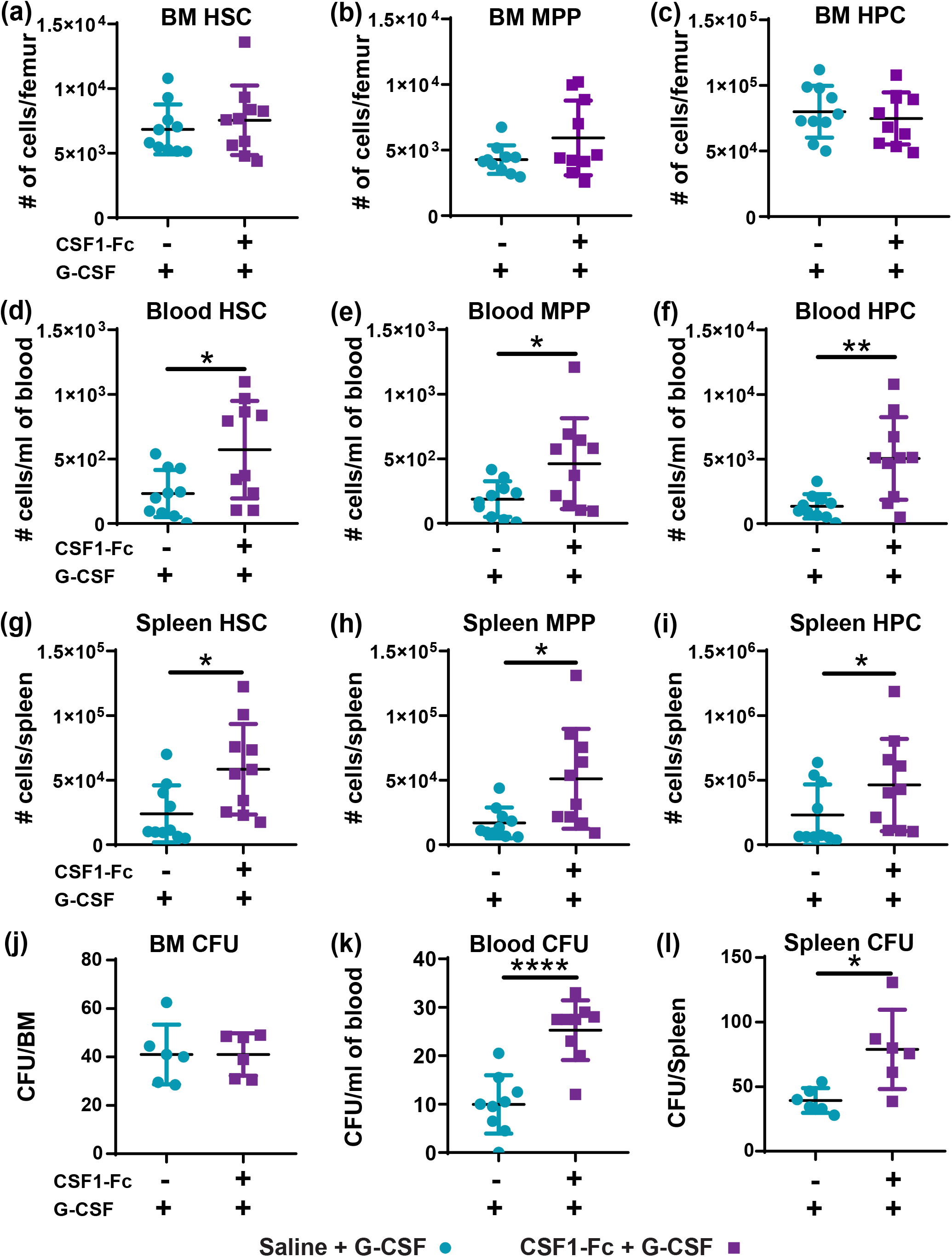
Sequential CSF1-Fc + G-CSF treatment mobilized HSPC more effectively than G-CSF alone. Experimental schematic is described in Fig. 5(a). **(a-i)** Flow cytometry analysis to enumerate number of **(a, d and g)** HSC, **(b, e and h)** MPP and **(c, f and i)** HPC in BM **(a-c)**, blood **(d-f)** and spleen **(g-i)** of mice treated as indicated. Population gating strategies are exemplified in Supp Fig. 4. Quantification of colony forming units (CFU) in the **(j)** BM (assay using single femur only), **(k)** blood and **(l)** spleen in mice treated as indicated. Each data point represents a separate mouse and bars are mean ± SD. Statistical analysis was performed using one-way ANOVA Tukey’s multiple comparison test where ****p<0.0001, **p<0.01 and *p<0.05.

The function of mobilised HSC from control and CSF1-Fc pre-treated mice was assessed using a competitive transplantation assay (Fig. 7a). Recipients that received a mobilised blood graft from G-CSF only control donors had detectable but low frequency of donor-derived blood leukocytes over 16 weeks post-transplant (Fig. 7b). Whereas the frequency of donor-derived blood leukocytes (Fig. 7b) of the donor graft was significantly increased when it was sourced from donors that had CSF1-Fc + G-CSF treatment. Calculation of repopulating units within the grafts showed a significant increase in the number of repopulating units/ml in the blood of mice mobilized with G-CSF plus CSF1-Fc compared to G-CSF alone (Fig. 7c). These results confirm that prior treatment with CSF1-Fc improved G-CSF-induced stem cell mobilization efficiency and reconstitution potential.

**Fig. 7.**
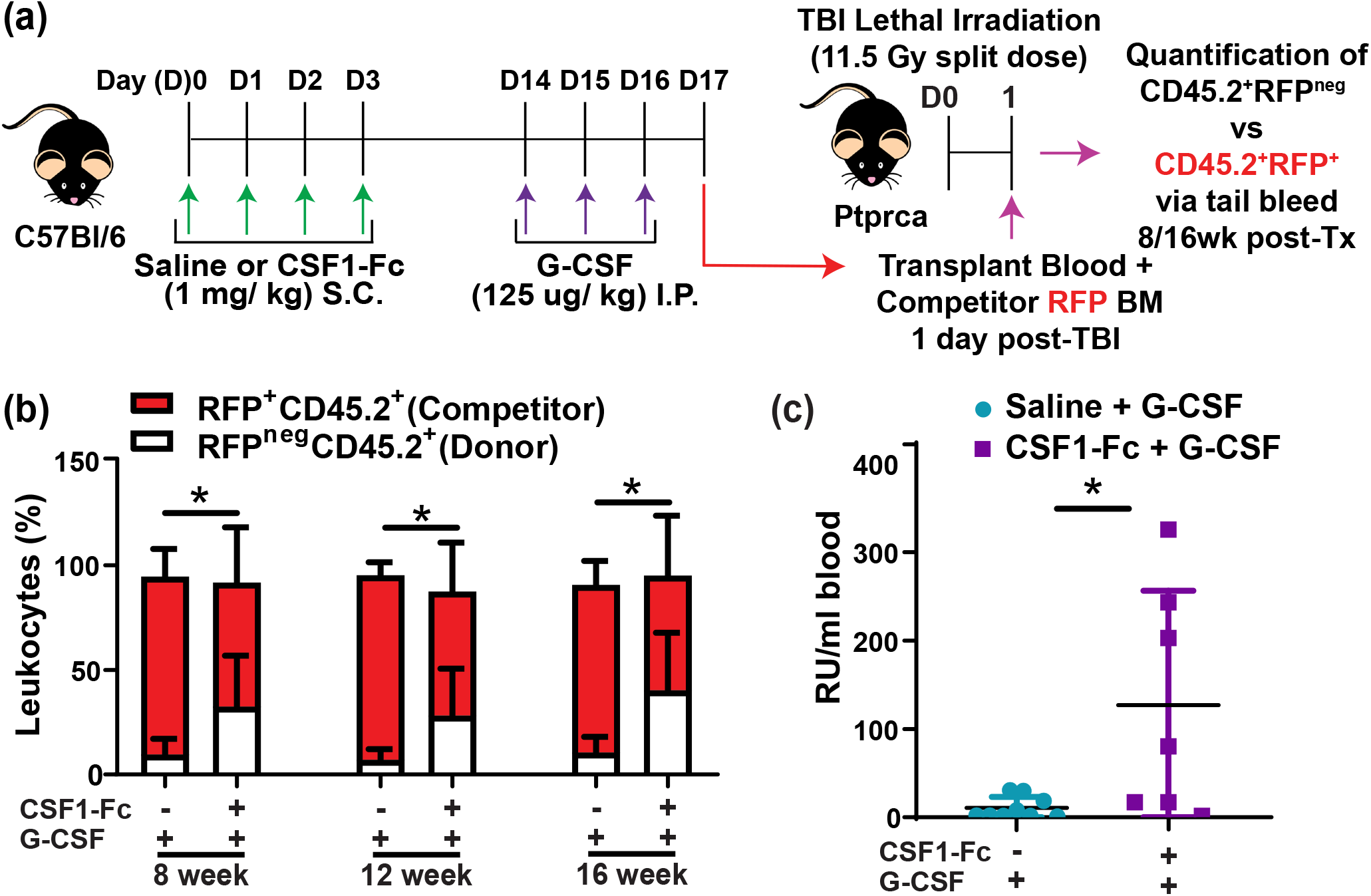
Combination of CSF1-Fc + G-CSF treatment improved the reconstitution potential of mobilized HSPC. **(a)** Schematic of competitive transplantation assay. Briefly, female donor C57BL/6 non-transgenic mice were treated with either once daily saline for 4 days followed by bi-daily G-CSF treatment (saline + G-CSF) 14 days later or once daily CSF1-Fc for 4 days followed by bi-daily G-CSF treatment (CSF1-Fc + saline) 14 days later as in Fig. 5a. The blood from donor C57BL/6 mice of two different treatment groups was then pooled with competitor BM from transgenic RFP mice at 17 days post initial CSF1-Fc treatment respectively and transplanted into lethally irradiated B6.SJL Ptprca recipients. Tail bleeds were performed at 8-, 12- and 16-weeks post-transplantation to determine chimerism. **(b)** Quantification of blood chimerism of RFP^neg^CD45.2^+^ donors (white bars) and RFP^+^CD45.2^+^ competitors (red bars) in recipient mice that were transplanted with blood from saline + G-CSF or CSF1-Fc + G-CSF treated donor mice. **(c)** Number of repopulating units (RU) per ml of blood transplanted in grafts collected from saline + G-CSF (light blue dots) and CSF1-Fc + G-CSF (purple squares) treated donors determined at 16 weeks post-competitive transplant. Data are mean ± SD. Evidence of data distribution non-normality was identified by the Kolmogorov–Smirnov test and statistical analysis was performed on data using by a Mann–Whitney U-test where **p<0.01 and *p<0.05.

## Discussion

The current study characterised hematopoietic impacts of treatment with a modified CSF1-Fc molecule that has improved drug qualities^15^. The expansion of monocytes and macrophages in BM and spleen following CSF1-Fc treatment was even more marked at day 7 (3 days after the last treatment) than at day 5 as studied previously^15^. This is consistent with the prolonged half-life of CSF1-Fc. Importantly, treatment impacts were rapidly reversible, reminiscent of transient effects of either 14 day continuous^49^ or daily^50^ infusion of unmodified CSF-1 in human clinical trials. At the time of peak response to CSF1-Fc, the prolonged excessive production of mature myeloid cells in BM and spleen occurred at the expense of normal haematopoiesis including disruption of BM HSC niche homeostasis. These impacts were again transient and largely resolved within a week of the peak treatment effect. We unexpectedly reveal that CSF1-Fc therapy caused a delayed increase in the total BM and spleen HSC pool and we showed that this could be manipulated to achieve enhanced HSC mobilisation for transplantation.

Both CSF1 therapeutic potential and elucidation of CSF1-mediated *in vivo* biology have been hampered due to the challenging practicalities associated with exogenous delivery of native CSF1. Continuous infusion of 150 µg/kg/day of native CSF1 in clinical trial resulted in only a transient increase in blood monocytes, peaking around day 7-8, despite ongoing growth factor infusion^49^ and a repeat high dose regimens did not result in cumulative effects in animal models^51,52^. Macrophages themselves clear CSF1 from the circulation by CSF1R-mediated endocytosis^53^. Consequently, the combination of treatment induced massive expansion of tissue macrophages, added to efficient renal clearance of native CSF1, likely culminate in rapid depletion of available growth factor from circulation despite ongoing treatment. Our and previous observations^15,19^ demonstrate that avoidance of renal clearance through use of modified CSF1-Fc is sufficient to achieve additive and sustained growth factor effects that parallel drug dose and predicted availability. Further preclinical application of CSF1-Fc has potential to accelerate discovery regarding the usefulness of targeting the CSF1-CSF1R axis to modulate macrophages in clinical applications including organ regeneration^54^, chemotherapy consolidation ^49^ and HSC transplantation^14,55^.

In spleen, resident macrophages can contribute to retention of HSC during extramedullary myelopoiesis^56^. Acute CSF1-Fc treatment expanded mature F4/80^+^ resident macrophages within spleen at the expense of lymphocyte, granulocyte and even monocyte frequency/retention. The expansion was skewed toward maturation of a subset of F4/80^+^CD169^+^ red pulp macrophages, that are only usually present at very low frequency in naïve spleen. Splenic erythropoiesis was significantly elevated at 1 week post-treatment and consequently at this time point, red pulp macrophages may have adapted to support extramedullary erythroid islands^32,57^. However, this unusual F4/80^+^CD169^+^ red pulp macrophage phenotype persisted even after splenic morphology was reinstated, extramedullary erythropoiesis had resolved, and macrophage frequency had returned to normal. We cannot determine in these experiments whether existing resident macrophages within spleen proliferated locally and changed phenotype or were replaced by recruited monocyte-derived macrophages. F4/80^+^CD169^+^ macrophages support HSC in BM^5,7^ and so we speculate that they acquire a similar function in spleen during resolution of CSF1-Fc treatment affects. Further studies are required to explicitly link the changed macrophage phenotype with improved HSC supportive function.

The potent monocytosis elicited by CSF1-Fc probably triggers known negative feedback loops directing compensatory reductions in lymphopoiesis and/or erythropoiesis^58^. We observed a significant impairment of BM B lymphopoiesis and temporary loss of lymphocytes in spleen. The observed reduction in BM B lymphopoiesis may be a secondary impact through altered osteoblast-lineage frequency or function. Osteoblast function and frequency can be influenced by both osteoclast and osteal macrophages^59^. CSF1-Fc treatment causes rapid expansion of bone-resorbing osteoclasts^15^ and osteal macrophages are CSF1-responsive^24,60^ and paradoxically systemic CSF1 treatment has an anabolic impact on bone^61^. Osteoblast-lineage cells in turn support B cell maturation^47^. Activation of this complex cellular feedback loop was not specifically examined in this study. The CSF1-Fc induced increase in monocyte/macrophages could also result in supraphysiologic accumulation of growth factors and cytokines that are expressed by these cells, many of which have the capacity to influence haematopoiesis. For example, excessive monopoiesis could result in elevated secretion of interleukin-1 and/or interferons, which are known to trigger HSC proliferation^62^. Further investigation is required to understand the secondary indirect impacts of CSF1-Fc treatment. A clinical challenge associated with autologous HSC transplantation is collection of sufficient HSC following mobilisation to achieve the required graft cell dose for successful transplant^1,2^. The increase of total available HSC pool induced by CSF1-Fc treatment presented herein could address this unmet need. Enhanced mobilisation of HSC into blood of mice treated with CSF1-Fc + G-CSF therapy was accompanied by increased CFU activity in blood. Importantly increased reconstitution of all blood lineages in recipient mice was demonstrated using grafts from combination therapy versus G-CSF alone. A remaining question is whether the long-term potential of the HSC within the CSF1-Fc + G-CSF donor graft is reduced due to CSF1-Fc exposure. While long-term serial transplant assays would be required to address this potential limitation, the collective data presented herein suggests that the stress induced on the HSPC and committed progenitor compartment is contained and reversible. Overall, this study has revealed the novel outcome of increased HSC pool as a consequence of CSF1-Fc therapy and that this might be a viable strategy to correct HSC availability deficits in patients at risk of poor mobilisation outcomes.

## Acknowledgments

RFP transgenic mouse line were generously provided by Professor Patrick Tam (Children’s Medical Research Institute) and Susie Nilsson (Monash University and CSIRO). Mr Kyle Williams (Mater Research Institute-The University of Queensland) provided technical assistance. Technical support was provided by the Translational Research Institute flow cytometry, histology, and microscopy core facilities and The University of Queensland Biological Resource’s Research Facility. The Translational Research Institute is supported by a grant from the Australian Government.

This work was supported by the National Health and Medical Research Council (project grants APP1102964 and APP1046590), the Mater Foundation, The University of Queensland International Scholarship (S.K.), an Australian Research Council Future Fellowship (ARP, FT150100335), and a National Health and Medical Research Council Senior Research Fellowship (JPL, APP1136130).

## Authorship Contributions

SK, AS and LJR: experimental design, performed experiments, data analysis, prepared figures and manuscript. SMM: experimental design, performed experiments, data analysis, prepared figures and approved manuscript. ACW, LB, MFC: performed experiments and approved manuscript. JPL and DAH: experimental design, data interpretation and manuscript editing. ARP: designed, led and coordinated the project, performed experiments, data analysis and figure and manuscript preparation.

## References

1. Perseghin P, Terruzzi E, Dassi M, et al. Management of poor peripheral blood stem cell mobilization: Incidence, predictive factors, alternative strategies and outcome. A retrospective analysis on 2177 patients from three major Italian institutions. Transfusion and Apheresis Science. 2009;41(1):33–37.

2. To LB, Levesque JP, Herbert KE. How I treat patients who mobilize hematopoietic stem cells poorly. Blood. 2011;118(17):4530–4540.

3. Domingues MJ, Nilsson SK, Cao B. New agents in HSC mobilization. Int J Hematol. 2017;105(2):141–152.

4. Winkler IG, Sims NA, Pettit AR, et al. Bone marrow macrophages maintain hematopoietic stem cell (HSC) niches and their depletion mobilizes HSCs. Blood. 2010;116(23):4815–4828.

5. Chow A, Lucas D, Hidalgo A, et al. Bone marrow CD169+ macrophages promote the retention of hematopoietic stem and progenitor cells in the mesenchymal stem cell niche. The Journal of Experimental Medicine. 2011;208(2):261–271.

6. Chang KH, Sengupta A, Nayak RC, et al. p62 is required for stem cell/progenitor retention through inhibition of IKK/NF-kappaB/Ccl4 signaling at the bone marrow macrophage-osteoblast niche. Cell Rep. 2014;9(6):2084–2097.

7. Kaur S, Raggatt LJ, Millard SM, et al. Self-repopulating recipient bone marrow resident macrophages promote long-term hematopoietic stem cell engraftment. Blood. 2018;132(7):735–749.

8. Duhrsen U, Villeval JL, Boyd J, Kannourakis G, Morstyn G, Metcalf D. Effects of recombinant human granulocyte colony-stimulating factor on hematopoietic progenitor cells in cancer patients. Blood. 1988;72(6):2074–2081.

9. Karpova D, Rettig MP, DiPersio JF. Mobilized peripheral blood: an updated perspective. F1000Res. 2019;8.

10. Tay J, Levesque JP, Winkler IG. Cellular players of hematopoietic stem cell mobilization in the bone marrow niche. Int J Hematol. 2017;105(2):129–140.

11. Christopher MJ, Rao M, Liu F, Woloszynek JR, Link DC. Expression of the G-CSF receptor in monocytic cells is sufficient to mediate hematopoietic progenitor mobilization by G-CSF in mice. The Journal of Experimental Medicine. 2011;208(2):251–260.

12. Hume DA, MacDonald KP. Therapeutic applications of macrophage colony-stimulating factor-1 (CSF-1) and antagonists of CSF-1 receptor (CSF-1R) signaling. Blood. 2012;119(8):1810–1820.

13. Hume DA, Caruso M, Ferrari-Cestari M, Summers KM, Pridans C, Irvine KM. Phenotypic impacts of CSF1R deficiencies in humans and model organisms. J Leukoc Biol. 2019.

14. Masaoka T, Shibata H, Ohno R, et al. Double-blind test of human urinary macrophage colony-stimulating factor for allogeneic and syngeneic bone marrow transplantation: effectiveness of treatment and 2-year follow-up for relapse of leukaemia. Br J Haematol. 1990;76(4):501–505.

15. Gow DJ, Sauter KA, Pridans C, et al. Characterisation of a novel Fc conjugate of macrophage colony-stimulating factor. Mol Ther. 2014;22(9):1580–1592.

16. Gow DJ, Garceau V, Kapetanovic R, et al. Cloning and expression of porcine Colony Stimulating Factor-1 (CSF-1) and Colony Stimulating Factor-1 Receptor (CSF-1R) and analysis of the species specificity of stimulation by CSF-1 and Interleukin 34. Cytokine. 2012;60(3):793–805.

17. Irvine KM, Caruso M, Cestari MF, et al. Analysis of the impact of CSF-1 administration in adult rats using a novel Csf1r-mApple reporter gene. J Leukoc Biol. 2019.

18. Sauter KA, Waddell LA, Lisowski ZM, et al. Macrophage colony-stimulating factor (CSF1) controls monocyte production and maturation and the steady-state size of the liver in pigs. Am J Physiol Gastrointest Liver Physiol. 2016;311(3):G533–547.

19. Pridans C, Raper A, Davis GM, et al. Pleiotropic Impacts of Macrophage and Microglial Deficiency on Development in Rats with Targeted Mutation of the Csf1r Locus. J Immunol. 2018;201(9):2683–2699.

20. Vintersten K, Monetti C, Gertsenstein M, et al. Mouse in red: red fluorescent protein expression in mouse ES cells, embryos, and adult animals. Genesis. 2004;40(4):241–246.

21. Levesque JP, Hendy J, Winkler IG, Takamatsu Y, Simmons PJ. Granulocyte colony-stimulating factor induces the release in the bone marrow of proteases that cleave c-KIT receptor (CD117) from the surface of hematopoietic progenitor cells. Exp Hematol. 2003;31(2):109–117.

22. Coquery CM, Loo W, Buszko M, Lannigan J, Erickson LD. Optimized protocol for the isolation of spleen-resident murine neutrophils. Cytometry A. 2012;81(9):806–814.

23. Irvine KM, Clouston AD, Gadd VL, et al. Deletion of Wntless in myeloid cells exacerbates liver fibrosis and the ductular reaction in chronic liver injury. Fibrogenesis & tissue repair. 2015;8:19–19.

24. Alexander KA, Chang MK, Maylin ER, et al. Osteal macrophages promote in vivo intramembranous bone healing in a mouse tibial injury model. J Bone Miner Res. 2011;26(7):1517–1532.

25. Forristal CE, Winkler IG, Nowlan B, Barbier V, Walkinshaw G, Levesque J-P. Pharmacologic stabilization of HIF-1α increases hematopoietic stem cell quiescence in vivo and accelerates blood recovery after severe irradiation. Blood. 2013;121(5):759–769.

26. Kwarteng EO, Heinonen KM. Competitive Transplants to Evaluate Hematopoietic Stem Cell Fitness. J Vis Exp. 2016(114).

27. Purton LE, Scadden DT. Limiting factors in murine hematopoietic stem cell assays. Cell Stem Cell. 2007;1(3):263–270.

28. Akashi K, Traver D, Miyamoto T, Weissman IL. A clonogenic common myeloid progenitor that gives rise to all myeloid lineages. Nature. 2000;404(6774):193–197.

29. Karsunky H, Inlay MA, Serwold T, Bhattacharya D, Weissman IL. Flk2+ common lymphoid progenitors possess equivalent differentiation potential for the B and T lineages. Blood. 2008;111(12):5562–5570.

30. Allman D, Pillai S. Peripheral B cell subsets. Curr Opin Immunol. 2008;20(2):149–157.

31. Mauri C, Bosma A. Immune regulatory function of B cells. Annu Rev Immunol. 2012;30:221–241.

32. Jacobsen RN, Forristal CE, Raggatt LJ, et al. Mobilization with granulocyte colony-stimulating factor blocks medullar erythropoiesis by depleting F4/80(+)VCAM1(+)CD169(+)ER-HR3(+)Ly6G(+) erythroid island macrophages in the mouse. Exp Hematol. 2014;42(7):547-561.e544.

33. Jacobsen RN, Nowlan B, Brunck ME, Barbier V, Winkler IG, Levesque JP. Fms-like tyrosine kinase 3 (Flt3) ligand depletes erythroid island macrophages and blocks medullar erythropoiesis in the mouse. Exp Hematol. 2016;44(3):207-212.e204.

34. Batoon L, Millard SM, Wullschleger ME, et al. CD169(+) macrophages are critical for osteoblast maintenance and promote intramembranous and endochondral ossification during bone repair. Biomaterials. 2017.

35. Inman CF, Rees LE, Barker E, Haverson K, Stokes CR, Bailey M. Validation of computer-assisted, pixel-based analysis of multiple-colour immunofluorescence histology. J Immunol Methods. 2005;302(1-2):156–167.

36. Levesque JP, Helwani FM, Winkler IG. The endosteal ‘osteoblastic’ niche and its role in hematopoietic stem cell homing and mobilization. Leukemia. 2010;24(12):1979–1992.

37. Borges da Silva H, Fonseca R, Pereira RM, Cassado Ados A, Alvarez JM, D’Imperio Lima MR. Splenic Macrophage Subsets and Their Function during Blood-Borne Infections. Front Immunol. 2015;6:480.

38. Perez OA, Yeung ST, Vera-Licona P, et al. CD169(+) macrophages orchestrate innate immune responses by regulating bacterial localization in the spleen. Sci Immunol. 2017;2(16).

39. Tay J, Bisht K, McGirr C, et al. Imaging flow cytometry reveals that G-CSF treatment causes loss of erythroblastic islands in the mouse bone marrow. Exp Hematol. 2020.

40. Seu KG, Papoin J, Fessler R, et al. Unraveling Macrophage Heterogeneity in Erythroblastic Islands. Front Immunol. 2017;8:1140.

41. Kiel MJ, Yilmaz ÖH, Iwashita T, Yilmaz OH, Terhorst C, Morrison SJ. SLAM Family Receptors Distinguish Hematopoietic Stem and Progenitor Cells and Reveal Endothelial Niches for Stem Cells. Cell. 2005;121(7):1109–1121.

42. Nestorowa S, Hamey FK, Pijuan Sala B, et al. A single-cell resolution map of mouse hematopoietic stem and progenitor cell differentiation. Blood. 2016;128(8):e20–31.

43. Hume DA, Freeman TC. Transcriptomic analysis of mononuclear phagocyte differentiation and activation. Immunol Rev. 2014;262(1):74–84.

44. Schmidl C, Renner K, Peter K, et al. Transcription and enhancer profiling in human monocyte subsets. Blood. 2014;123(17):e90–99.

45. Sasmono RT, Ehrnsperger A, Cronau SL, et al. Mouse neutrophilic granulocytes express mRNA encoding the macrophage colony-stimulating factor receptor (CSF-1R) as well as many other macrophage-specific transcripts and can transdifferentiate into macrophages in vitro in response to CSF-1. J Leukoc Biol. 2007;82(1):111–123.

46. Shen H, Yu H, Liang PH, et al. An acute negative bystander effect of gamma-irradiated recipients on transplanted hematopoietic stem cells. Blood. 2012;119(15):3629–3637.

47. Winkler IG, Bendall LJ, Forristal CE, et al. B-lymphopoiesis is stopped by mobilizing doses of G-CSF and is rescued by overexpression of the anti-apoptotic protein Bcl2. Haematologica. 2013;98(3):325–333.

48. Carsetti R. The development of B cells in the bone marrow is controlled by the balance between cell-autonomous mechanisms and signals from the microenvironment. J Exp Med. 2000;191(1):5–8.

49. Cole DJ, Sanda MG, Yang JC, et al. Phase I trial of recombinant human macrophage colony-stimulating factor administered by continuous intravenous infusion in patients with metastatic cancer. J Natl Cancer Inst. 1994;86(1):39–45.

50. Ohno R, Miyawaki S, Hatake K, et al. Human urinary macrophage colony-stimulating factor reduces the incidence and duration of febrile neutropenia and shortens the period required to finish three courses of intensive consolidation therapy in acute myeloid leukemia: a double-blind controlled study. J Clin Oncol. 1997;15(8):2954–2965.

51. Lloyd SA, Simske SJ, Bogren LK, Olesiak SE, Bateman TA, Ferguson VL. Effects of combined insulin-like growth factor 1 and macrophage colony-stimulating factor on the skeletal properties of mice. In Vivo. 2011;25(3):297–305.

52. Kodama H, Yamasaki A, Nose M, et al. Congenital osteoclast deficiency in osteopetrotic (op/op) mice is cured by injections of macrophage colony-stimulating factor. J Exp Med. 1991;173(1):269–272.

53. Jenkins SJ, Hume DA. Homeostasis in the mononuclear phagocyte system. Trends Immunol. 2014;35(8):358–367.

54. Stutchfield BM, Antoine DJ, Mackinnon AC, et al. CSF1 Restores Innate Immunity After Liver Injury in Mice and Serum Levels Indicate Outcomes of Patients With Acute Liver Failure. Gastroenterology. 2015;149(7):1896–1909 e1814.

55. Kandalla PK, Sarrazin S, Molawi K, et al. M-CSF improves protection against bacterial and fungal infections after hematopoietic stem/progenitor cell transplantation. J Exp Med. 2016;213(11):2269–2279.

56. Dutta P, Hoyer FF, Grigoryeva LS, et al. Macrophages retain hematopoietic stem cells in the spleen via VCAM-1. The Journal of Experimental Medicine. 2015;212(4):497–512.

57. Chow A, Huggins M, Ahmed J, et al. CD169(+) macrophages provide a niche promoting erythropoiesis under homeostasis and stress. Nat Med. 2013;19(4):429–436.

58. Zhao JL, Baltimore D. Regulation of stress-induced hematopoiesis. Curr Opin Hematol. 2015;22(4):286–292.

59. Pettit AR, Chang MK, Hume DA, Raggatt LJ. Osteal macrophages: a new twist on coupling during bone dynamics. Bone. 2008;43(6):976–982.

60. Raggatt LJ, Wullschleger ME, Alexander KA, et al. Fracture healing via periosteal callus formation requires macrophages for both initiation and progression of early endochondral ossification. Am J Pathol. 2014;184(12):3192–3204.

61. Lloyd SA, Yuan YY, Simske SJ, Riffle SE, Ferguson VL, Bateman TA. Administration of high-dose macrophage colony-stimulating factor increases bone turnover and trabecular volume fraction. J Bone Miner Metab. 2009;27(5):546–554.

62. Pietras EM. Inflammation: a key regulator of hematopoietic stem cell fate in health and disease. Blood. 2017;130(15):1693–1698.

