## Supplemental Figures and Tables for "Stable colony stimulating factor 1 fusion protein treatment increases HSC pool and enhances their mobilisation in mice"

**Supplementary Data: Systemic treatment with a stable colony stimulating factor 1 fusion protein increases total hematopoietic stem cells and enhances their subsequent mobilisation in mice.**

**Supp. Fig. 1. Myeloid lineage gating strategy in BM and spleen of saline or CSF1-Fc treated mice.**

Representative flow cytometry plots of myeloid lineage in saline and CSF1-Fc treated mice at day (D)7 and D14 post-treatment. (a) BM monocytes were first gated as F4/80$^{+}$Ly6G$^{neg}$ cells (orange gate) then further gated as VCAM-1$^{neg}$CD115$^{+}$ cells (purple gate) that were predominantly CD11b$^{+}$ (not shown). (b) BM granulocytes were identified as CD11b$^{+}$Ly6G$^{+}$ cells (green gate). (c) Splenic monocytes were first gated as F4/80$^{+}$Ly6G$^{neg}$ cells (orange gate) and then further gated as VCAM-1$^{neg}$CD115$^{+}$ cells (purple gate) that were predominantly CD11b$^{+}$ (not shown). (d) Splenic granulocytes were identified as CD11b$^{+}$Ly6G$^{+}$ cells (green gate).

**Supp. Fig. 2. Lymphocyte lineage gating strategy in BM and spleen of saline or CSF1-Fc treated mice.**

Representative flow cytometry plots of B cell subsets and mature T cells in saline and CSF1-Fc treated mice at day (D)7 and D14 post-treatment in BM (a) and spleen (b). (a) In BM CD11b$^{neg}$ cells were further gated as CD45R(B220)$^{+}$CD3$^{neg}$ B cells (green gate) and CD45R(B220)$^{neg}$CD3$^{+}$ T cells (red gate). Total CD45R(B220)$^{+}$ BM cells were analysed for IgM and IgD analysis where Ig$^{neg}$ B cells (orange gate) were further profiled for CD19 and CD43 expression to determine CD19$^{neg}$CD43$^{+}$ Pre-pro-B cells (blue gate), CD19$^{+}$CD43$^{+}$ Pro-B cells (green gate) and CD19$^{+}$CD43$^{neg}$ Pre-B cells (purple gate). (b) Spleens were analysed for the gated fractions CD11b$^{-}$CD3$^{neg}$ cells (orange gate) and CD11b$^{-}$CD3$^{+}$ T cells (red gate).

CD11b$^{neg}$CD3$^{neg}$ cells were further profiled for total CD4R(B220)$^{+}$ B cells (green gate). The CD4R(B220)$^{+}$CD93$^{+}$ fraction (orange gate) were further analysed to identify transitional B cells: IgM$^{+}$CD23$^{neg}$ transitional-1 (T1) B cells (purple gate), IgM$^{+}$CD23$^{+}$ T2 B cells (green gate) and IgM$^{neg}$CD23$^{+}$ T3 B cells (blue gate).

**Supp. Fig. 3. Immunofluorescence analysis of splenic architecture disruption and CD169$^{+}$ cell frequency in white pulp 14 days post-CSF1-Fc treatment.**

**(a)** Immunofluorescence labelling of CD169 (Red) and CD3 (Green) in spleen sections of mice treated with saline (top panel) or CSF1-Fc at 7 days (D7; middle panel) and 14 days (D14; bottom panel) post-first injection. Magnification = 60X; scale bar = 500 µm. Inset magnification = 600X. **(b-c)** Morphometric analysis of **(b)** percent area CD3 immunolabelling and **(c)** CD169 immunolabelling within T cell zones in spleens of saline controls or CSF1-Fc treated mice at the D7 and D14 time points. Each data point represents a separate mouse and bars are mean ± SD. Evidence of data distribution non-normality was identified by the Kolmogorov–Smirnov test and statistical analysis was performed on data using by a Mann–Whitney U-test where ****p < 0.0001.

**Supp. Fig. 4. HSPC gating strategy in BM, spleen and liver of saline or CSF1-Fc treated mice.**

Representative flow cytometry analysis for committed progenitors and HSPC subsets in saline (top panel) or day (D) 7 (middle panel) and D14 (bottom panel) CSF1-Fc treated mice in (a) BM, (b) spleen and (c) liver. Committed progenitor cells were gated as lineage negative, c-Kit$^{+}$ and Sca1$^{neg}$ cells (black gate). HSPC were first gated as LSK (lineage negative, c-Kit$^{+}$ and Sca1$^{+}$ cells, orange gate). These were further fractioned as: CD48$^{neg}$CD150$^{+}$ HSC (blue gate), CD48$^{neg}$CD150$^{neg}$ MPP (green gate) and CD48$^{+}$ HPC (purple gate).

**Supp. Fig. 5. Competitive transplantation model and quantification of leukocyte chimerism in blood confirmed CSF1-Fc reduced BM HSC repopulation potential at 7 days after treatment.**

**(a)** Schematic of competitive transplantation assay. Briefly, female donor C57BL/6 non-transgenic mice were treated with either daily saline or CSF1-Fc for 4 days. The donor C57BL/6 BM of the two different treatment groups was collected 7 days post-first CSF1-Fc treatment and pooled with competitor UBG-GFP BM prior to transplant into lethally irradiated C57BL/6 non-transgenic recipients. (**b**) Tail bleeds were performed at 8-, 12- and 16-week post-transplantation and analysed for blood chimerism of CD45.2$^{+}$GFP$^{neg}$ donors (white bars) and CD45.2$^{+}$GFP$^{+}$ competitors (green bars) in recipient mice that receive BM from saline or CSF1-Fc treated donor mice. Data are mean ± SD. Statistical analysis was performed using one-way ANOVA Tukey's multiple comparison test where *$p<0.05$, n = 8 to 10 mice/group.

**Supp. Fig. 6. Myeloid and lymphoid progenitor cell gating strategy in BM, spleen and liver of saline or CSF1-Fc treated mice.**

Representative flow cytometry analysis for myeloid progenitor cells in (a) BM (b) spleen and (c) liver and lymphoid progenitor cells in (d) BM, (e) spleen and (f) liver in saline or day (D) 7 and D14 CSF1-Fc treated mice. (a-c) For myeloid progenitor cells, Lineage$^{neg}$Sca-1$^{neg}$cKit$^{+}$ cells (orange gate) were gated into CD16/CD32$^{int}$CD34$^{+}$ common myeloid progenitors (CMP; blue gate), CD16/CD32$^{+}$CD34$^{+}$ granulocyte-macrophage progenitors (GMP; purple gate) and CD16/CD32$^{neg}$CD34$^{neg}$ megakaryocyte erythroid progenitors (MEP; green gate). (d-f) For lymphoid progenitor cells, Lineage$^{neg}$IL-7rα$^{+}$ (orange gate) were gated into Sca-1$^{int}$c-Kit$^{int}$ cells (purple gate) and the expression of Flk2 (receptor for Flt3 ligand) was then analysed on this population of cells to determine numbers of common lymphoid progenitors (CLP; green gate).

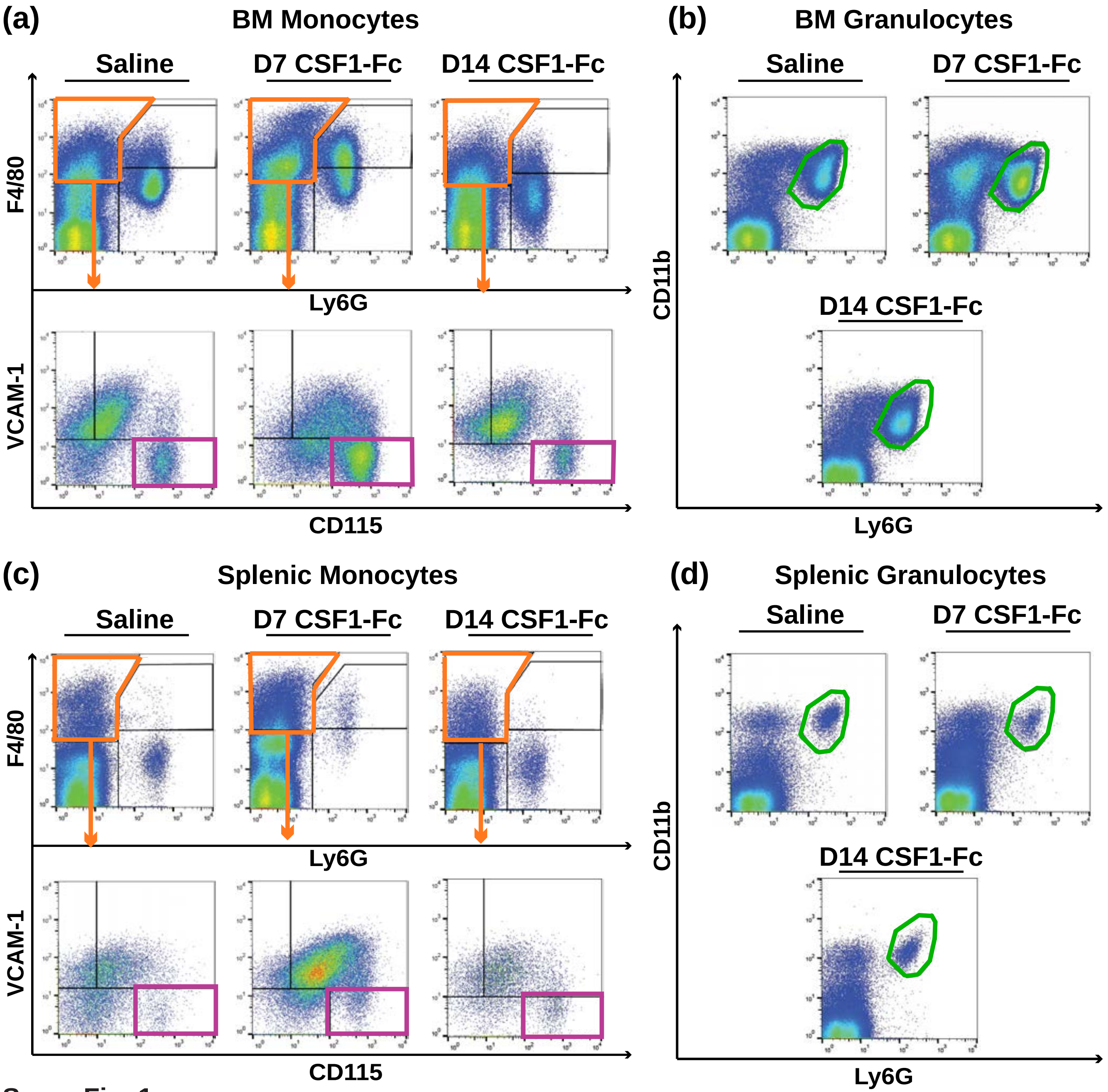
(a)
BM Monocytes
Saline
D7 CSF1-Fc
D14 CSF1-Fc
F4/80
Ly6G
VCAM-1
CD115
(b)
BM Granulocytes
Saline
D7 CSF1-Fc
D14 CSF1-Fc
CD11b
Ly6G
(c)
Splenic Monocytes
Saline
D7 CSF1-Fc
D14 CSF1-Fc
F4/80
Ly6G
VCAM-1
CD115
(d)
Splenic Granulocytes
Saline
D7 CSF1-Fc
D14 CSF1-Fc
CD11b
Ly6G

Supp. Fig. 1

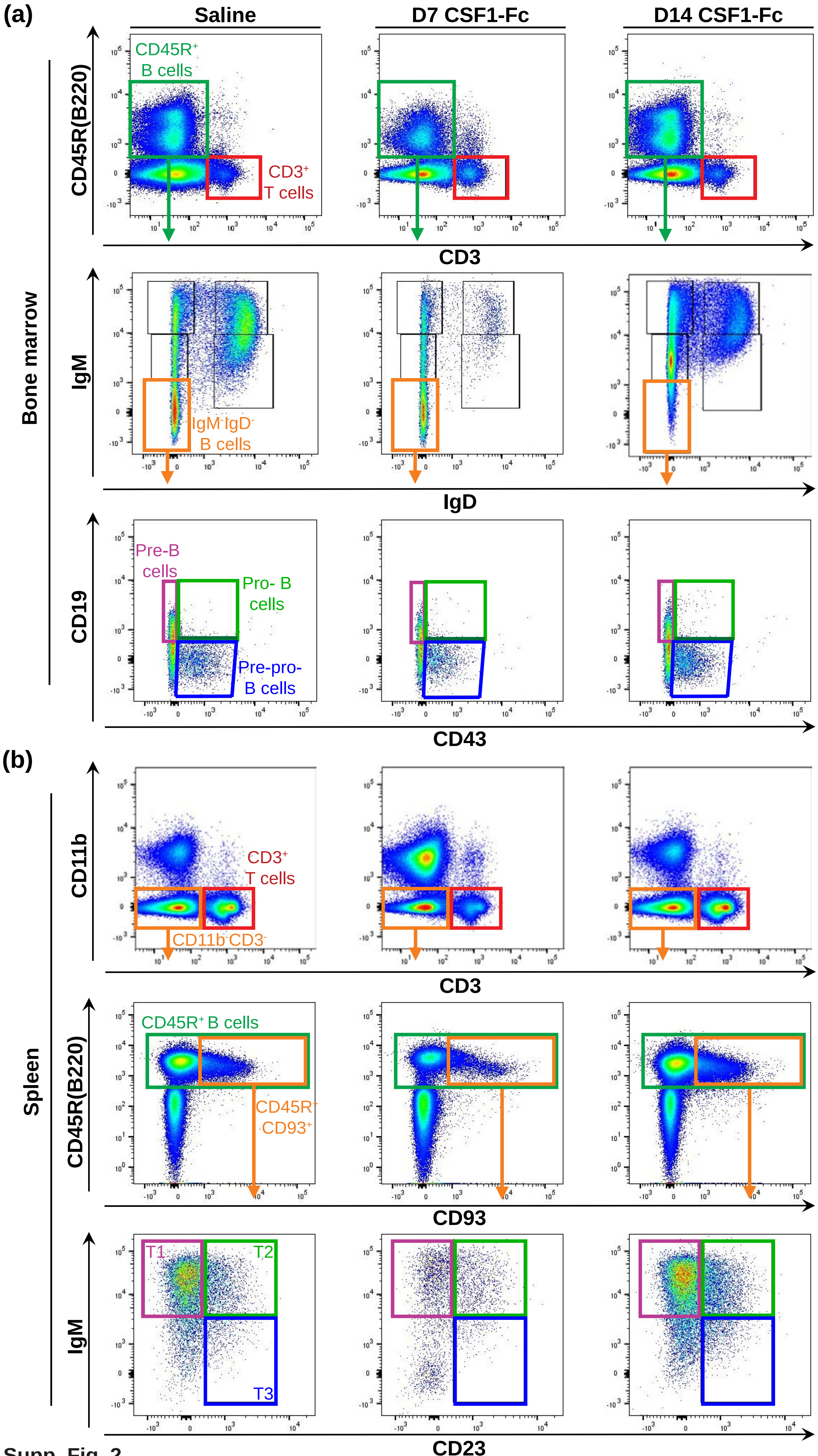
(a)
Saline
D7 CSF1-Fc
D14 CSF1-Fc
Bone marrow
CD45R(B220)
CD45R+ B cells
CD3+ T cells
CD3
IgM
IgM-IgD- B cells
IgD
CD19
Pre-B cells
Pro- B cells
Pre-pro- B cells
CD43
(b)
Spleen
CD11b
CD3+ T cells
CD11b-CD3-
CD3
CD45R(B220)
CD45R+ B cells
CD45R+ CD93+
CD93
IgM
T1
T2
T3
CD23

Supp. Fig. 2

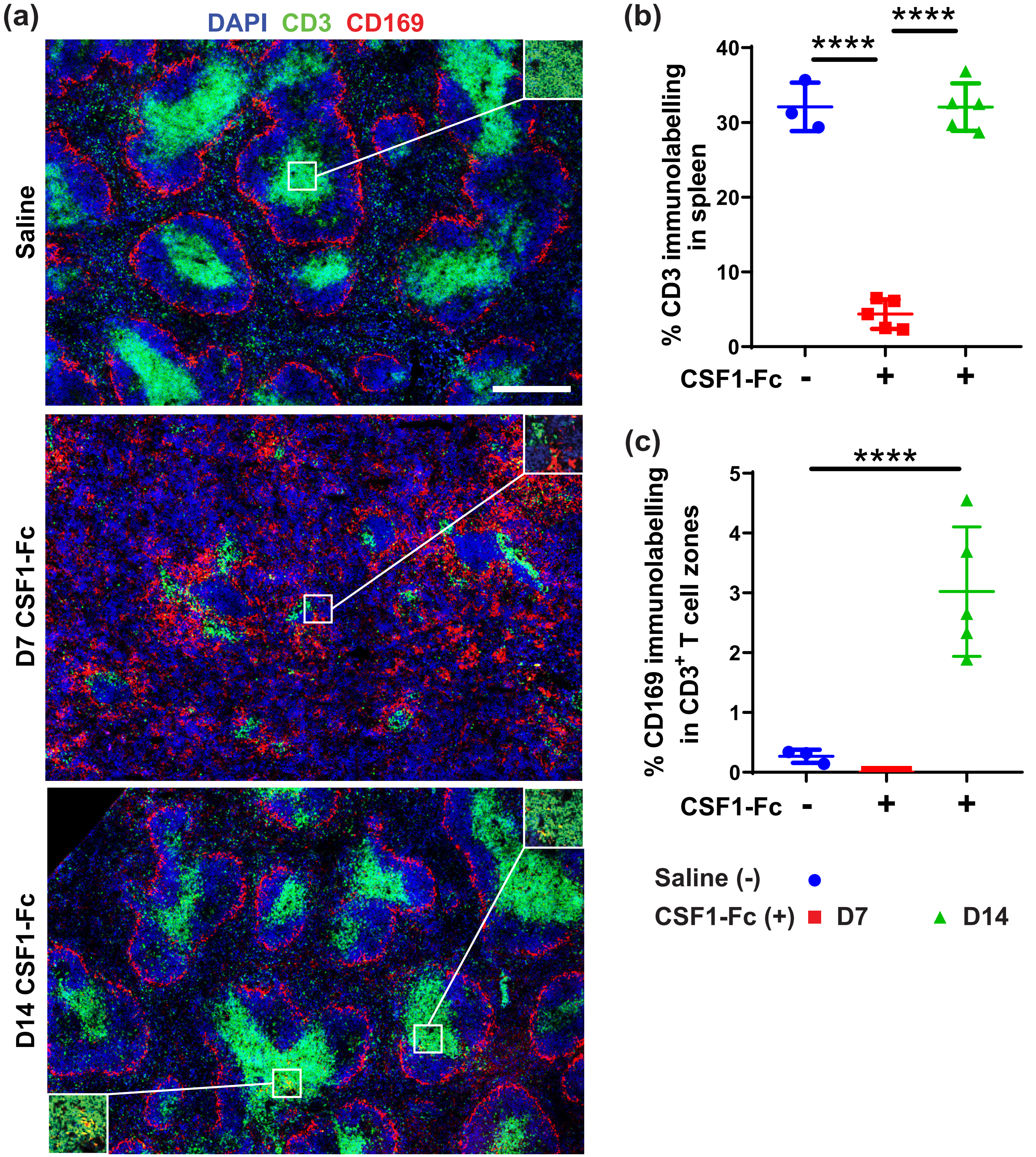
(a)
DAPI CD3 CD169
Saline
D7 CSF1-Fc
D14 CSF1-Fc
(b)
% CD3 immunolabelling in spleen
40
30
20
10
0
****
****
CSF1-Fc - + +
(c)
% CD169 immunolabelling in $CD3^+$ T cell zones
5
4
3
2
1
0
****
CSF1-Fc - + +
Saline (-)
CSF1-Fc (+) D7 D14

Supp. Fig. 3

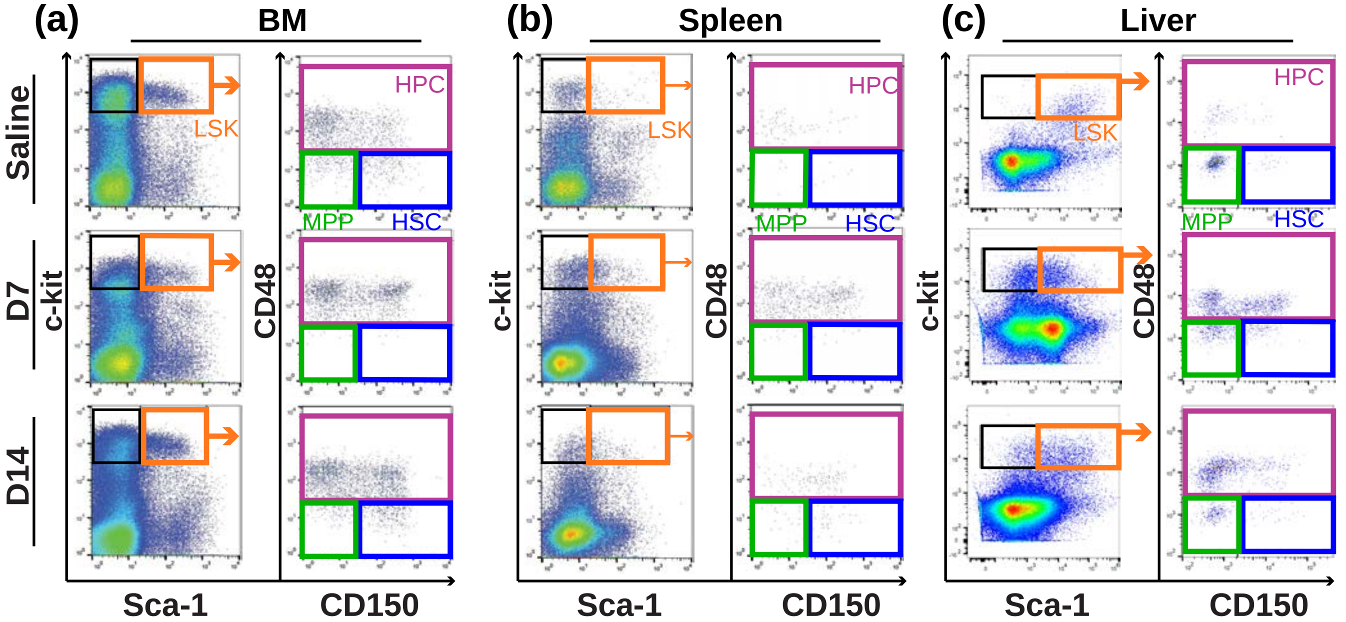
(a)
BM
(b)
Spleen
(c)
Liver
Saline
D7
D14
c-kit
CD48
LSK
HPC
MPP
HSC
Sca-1
CD150

Supp. Fig. 4

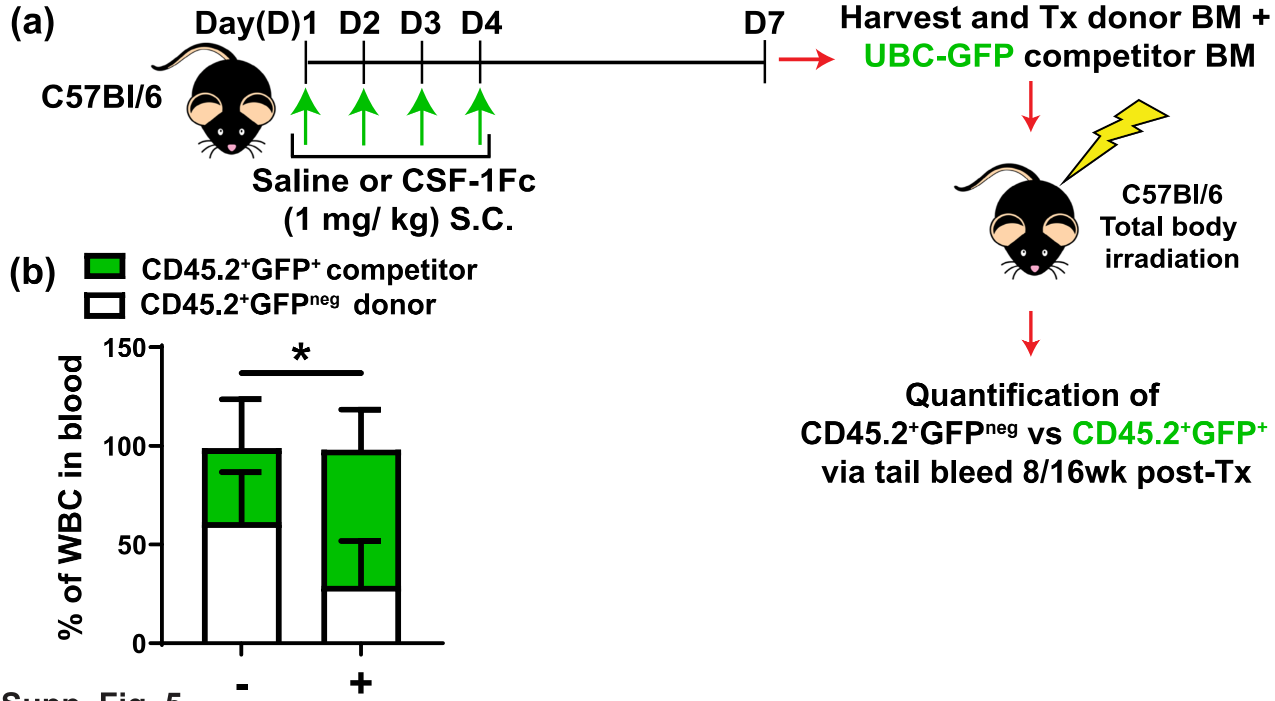
(a)
Day(D)1 D2 D3 D4 D7
C57Bl/6
Saline or CSF-1Fc
(1 mg/ kg) S.C.
Harvest and Tx donor BM +
UBC-GFP competitor BM
C57Bl/6
Total body
irradiation
Quantification of
CD45.2+GFPneg vs CD45.2+GFP+
via tail bleed 8/16wk post-Tx
(b)
CD45.2+GFP+ competitor
CD45.2+GFPneg donor
% of WBC in blood
150
100
50
0
*
-
+

Supp. Fig. 5

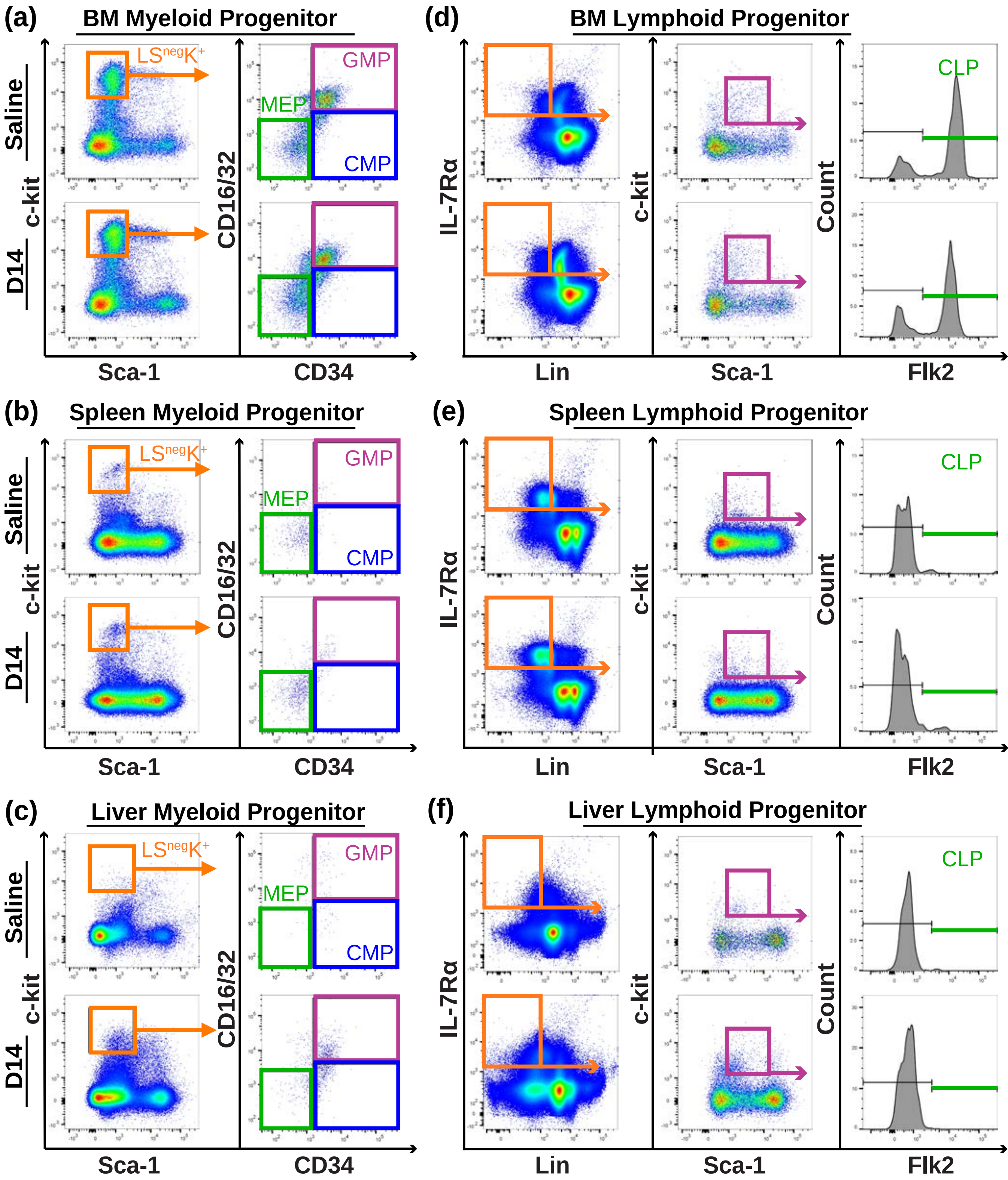
(a)
BM Myeloid Progenitor
LSnegK+
GMP
MEP
CMP
Saline
D14
c-kit
CD16/32
Sca-1
CD34
(d)
BM Lymphoid Progenitor
CLP
IL-7Rα
c-kit
Count
Lin
Sca-1
Flk2
(b)
Spleen Myeloid Progenitor
(e)
Spleen Lymphoid Progenitor
(c)
Liver Myeloid Progenitor
(f)
Liver Lymphoid Progenitor

Supp. Fig. 6

**Supplementary Table 1. Flow cytometry antibody details.**

| Panel/Tissues | Specificity | Flourochrome | Clone | Supplier | Catalogue Number |
|---|---|---|---|---|---|
| **Myeloid panel** | Ly6G | BV785 | 1A8 | Biolegend® | 127645 |
| **All tissues** | CD115 | BV605 | AFS98 | Biolegend® | 135517 |
| | CD11b | BV510 | M1/70 | Biolegend® | 101245 |
| | VCAM-1 | BV421 | 51-10C9 | BD Biosciences | 744309 |
| | CD169 | PE-Cy7 | 3D6.112 | Biolegend® | 142411 |
| | F4/80 | AF647 | BM8 | Biolegend® | 123121 |
| **Lymphocyte Panel** | IgD | BUV395 | 11-26c.2a | BD Biosciences | 564274 |
| **All tissues** | CD93 | BV650 | AA4.1 | BD Biosciences | 563807 |
| | IgM | BV605 | RMM-1 | Biolegend® | 406523 |
| | CD11b | BV510 | M1/70 | Biolegend® | 101245 |
| | CD4 | PB | GK1.5 | Biolegend® | 100427 |
| | CD3 | FITC | 17A2 | Biolegend® | 100203 |
| | CD19 | PerCP-CY5.5 | 1D3/CD19 | Biolegend® | 152405 |
| | CD43 | PE | S11 | Biolegend® | 143205 |
| | CD8a | PE-Cy7 | 53-6.7 | Biolegend® | 100721 |
| | CD23 | AF700 | B3B4 | Biolegend® | 101631 |
| | CD45R(B220) | APC-Cy7 | RA3-6B2 | Biolegend® | 103223 |
| **LSK Panel** | CD3 | FITC | 17A2 | Biolegend® | 100203 |
| **All tissues** | CD5 | FITC | 53-7.3 | Biolegend® | 100605 |
| | CD11b | FITC | M1/70 | Biolegend® | 101205 |
| | CD45R(B220) | FITC | RA3-6B2 | BD Biosciences | 553087 |
| | GR-1 | FITC | RB6-8C5 | Biolegend® | 108405 |
| | TER119 | FITC | TER-119 | Biolegend® | 116205 |

| **Panel/Tissues** | **Specificity** | **Flourochrome** | **Clone** | **Supplier** | **Catalogue Number** |
|---|---|---|---|---|---|
| | CD150 | BV650 | TC15-12F12.2 | Biolegend® | 115931 |
| | CD48 | PB | HM48-1 | Biolegend® | 103417 |
| | Sca-1 | Pe-Cy7 | E13-161.7 | Biolegend® | 122513 |
| | cKit | APC | 2B8 | Biolegend® | 105811 |
| **Myeloid and Lymphoid Progenitors Panel** | IL-7Ra | BV785 | A7R34 | Biolegend® | 135037 |
| | Sca-1 | BV510 | D7 | Biolegend® | 108129 |
| **All tissues** | CD3 | PB | 17A2 | Biolegend® | 100213 |
| | CD5 | PB | 53-7.3 | Biolegend® | 100641 |
| | CD11b | PB | M1/70 | Biolegend® | 101223 |
| | CD45R(B220) | PB | RA3-6B2 | Biolegend® | 103230 |
| | GR-1 | PB | RB6-8C5 | Biolegend® | 108429 |
| | TER119 | PB | TER-119 | Biolegend® | 116231 |
| | CD16/32 | PE | 93 | Biolegend® | 101301 |
| | CD34 | e660 | RAM34 | Thermo Fisher Scientific | 50-0341-82 |
| | cKit | APC-Cy7 | 2B8 | | 105825 |
| **Erythroblast Lineage Panel** | Hoechst 33342 | N/A | N/A | Thermo Fisher Scientific | H3570 |
| | CD71 | PE | RI7217 | Biolegend® | 113807 |
| | TER119 | PE-Cy7 | TER-119 | Biolegend® | 116221 |
| | CD45 | APC | 30-F11 | Biolegend® | 103111 |
| **Viability Dye** | 7-Aminoactinomycin D (7AAD) | N/A | N/A | Thermo Fisher Scientific | A1310 |
